# Aberrant PCM1 accumulation in Trisomy 21 mislocalizes E3 ligases, delaying primary ciliogenesis

**DOI:** 10.64898/2026.09.14.751618

**Authors:** Bailey L. McCurdy, Chad G. Pearson

**Affiliations:** Department of Cell and Developmental Biology, University of Colorado School of Medicine, Aurora, CO 80045-2537; Linda Crnic Institute for Down Syndrome, University of Colorado School of Medicine, Aurora, CO 80045-2537

**Keywords:** Primary ciliogenesis, Centriolar uncapping, Centrosome, Pericentrosome, Trisomy 21, Centriolar satellites

## Abstract

Primary cilia are microtubule-based extracellular signaling structures essential for development and tissue homeostasis, and their defects can cause ciliopathies. Trisomy 21, the cause of Down syndrome, also disrupts cilia formation and function. Here we show that Pericentrin (PCNT), a chromosome 21 resident gene whose protein is elevated in Trisomy 21, impairs primary ciliogenesis by delaying mother centriole uncapping. Elevated PCNT forms pericentrosomal assemblies that promote the accumulation of PCM1 in the pericentrosomal compartment. PCM1 binds and localizes the CP110 E3 ubiquitin ligases HERC2 and MIB1 to the centrosome, so mislocalizing PCM1 depletes these ligases from the centrosome, lowering CP110 ubiquitination and delaying uncapping. Reducing PCNT rescues these defects, whereas elevating PCM1 phenocopies them, establishing PCM1 accumulation as a critical mediator of uncapping. These findings reveal how a dosage-sensitive chromosome 21 gene disrupts ciliogenesis and show that centrosome function depends not only on PCM protein composition but on the spatial partitioning of proteins between the centrosome and pericentrosomal compartments.

**SUMMARY:** Elevated Pericentrin in Trisomy 21 remodels the pericentrosomal space, causing PCM1-dependent sequestration of CP110 ubiquitin ligases away from the centrosome. This delays CP110 degradation, centriolar uncapping, and primary ciliogenesis revealing a mechanism linking centrosome architecture to cilium assembly.

## INTRODUCTION

Primary cilia project from the cell surface and mediate signaling between cells. Cilia-dependent signaling pathways are essential for development and tissue homeostasis, and mutations in genes affecting cilia result in a collection of diseases known as ciliopathies (Bisgrove & Yost, 2006; Goetz & Anderson, 2010). Trisomy 21 (T21), the cause of Down syndrome, results from supernumerary copy number of human chromosome 21 (Hattori et al., 2000). Notably, individuals with Down syndrome display phenotypic overlap with individuals with ciliopathies, and T21 itself causes defects in primary cilia formation and signaling (Currier et al., 2012; Galati et al., 2018; Jewett et al., 2023; McCurdy et al., 2022; Ripoll et al., 2012; Roper et al., 2006).

Defective primary ciliogenesis in T21 is caused, at least in part, by overexpression of Pericentrin (PCNT), a chromosome 21 encoded component of the pericentriolar material (PCM) (Galati et al., 2018; Jewett et al., 2023; McCurdy et al., 2022). T21 elevates PCNT, and as a PCM scaffold protein, PCNT recruits PCM components and promotes microtubule (MT) nucleation from the centrosome, the microtubule organizing center (MTOC) formed by two centrioles surrounded by the PCM (Bettencourt-Dias et al., 2011; Dictenberg et al., 1998; Doxsey et al., 1994; Fong et al., 2008; Gavilan et al., 2018). PCM composition and abundance are tightly regulated during the cell cycle: the PCM expands during mitosis to support spindle MT nucleation and organization, then it is reduced during G0/G1 to facilitate efficient MT-dependent transport of components required for primary ciliogenesis to the mother centriole, from which the cilium nucleates (Conduit et al., 2015; Lee & Rhee, 2011; McCurdy et al., 2022). The formation of primary cilia during quiescence, a period of reduced PCM abundance, is particularly susceptible to disruption by excess PCNT. The mechanistic impact of elevated PCNT on primary ciliogenesis remains unknown.

In T21, elevated PCNT protein accumulates in cytoplasmic assemblies surrounding centrosomes, a region called the pericentrosome (Dammermann & Merdes, 2002; Galati et al., 2018; Jewett et al., 2023; McCurdy et al., 2022). These assemblies recruit y-tubulin and nucleate pericentrosomal MTs, resulting in mislocalized proteins required for ciliogenesis, including the centriolar satellite protein PCM1 (Jewett et al., 2023; McCurdy et al., 2022). Centriolar satellites are discrete pericentrosomal and cytoplasmic structures that interact with ciliogenesis-associated proteins to promote primary cilium assembly, though the mechanisms governing their composition, function, and dynamics at ciliogenesis onset remain poorly understood (Bergström et al., 2016; Hall et al., 2023; Kubo et al., 1999; Odabasi et al., 2019; Prosser & Pelletier, 2020; Tollenaere et al., 2015). One centriolar satellite protein, PCM1, is required for ciliogenesis in some cell types but its diverse functions remain poorly understood (Hall et al., 2023; Odabasi et al., 2019; Prosser & Pelletier, 2020; Tollenaere et al., 2015; Wang et al., 2016). In T21, PCNT assemblies colocalize with PCM1 in the pericentrosomal region, suggesting that excess PCNT sequesters PCM1 and elevates its local concentration in the pericentrosomal region (McCurdy et al., 2022). Whether this mislocalization or aberrant accumulation of PCM1 impairs the onset of primary ciliogenesis remains to be determined, and addressing this question may clarify both the mechanisms for ciliogenesis defects caused by T21 and the role of PCM1 in primary cilium assembly.

The onset of primary cilia assembly is governed at the mother centriole, where removal of the CP110/CEP97 cap licenses ciliary axoneme formation (Sorokin, 1962; Spektor et al., 2007; Tsang et al., 2008; Yadav et al., 2016). The mother centriole possesses appendages that nucleate MTs, recruit preciliary vesicles, and facilitate plasma membrane docking, and upon conversion to a basal body it nucleates ciliary axoneme formation (Bettencourt-Dias & Glover, 2007; Kumar & Reiter, 2021; Shakya & Westlake, 2021; Tanos et al., 2013). Ciliary axoneme formation requires removal of the CP110/CEP97 cap, which otherwise blocks axoneme elongation (Iyer et al., 2025; Lu et al., 2026; Zhao et al., 2023). CP110 also functions in centriole duplication and maintenance of centriole length during the cell cycle, and these diverse roles for CP110 protein are controlled largely by CP110 protein levels. Loss of CP110 leads to elongated centrioles and aberrant primary cilia formation in proliferating cells, whereas excess CP110 suppresses primary ciliogenesis in quiescent cells (Iyer et al., 2025; Kleylein-Sohn et al., 2007; Schmidt et al., 2009; Spektor et al., 2007). Therefore, CP110 levels must be tuned precisely at ciliogenesis onset. This is achieved through proteasome-mediated degradation driven by E3 ubiquitin ligases MIB1 and HERC2, which localize to the centrosome, ubiquitinate CP110, and promote degradation of the CP110/CEP97 cap (Xie et al., 2023). Critically, PCM1 interacts with and facilitates MIB1 and HERC2 localization to the centrosome (Xie et al., 2023). PCM1 mislocalization, as occurs in T21, is a candidate mechanism by which centrosomal MIB1 and HERC2 are reduced, impairing CP110 uncapping and primary ciliogenesis.

Here, we discover a CP110 uncapping delay in T21. Excess PCNT drives pericentrosomal accumulation of PCM1, which sequesters MIB1 and HERC2 and reduces their centrosomal levels, attenuating CP110 ubiquitination and uncapping. Overexpression of PCM1 alone recapitulates this phenotype. Directing PCM1 to the cell periphery depletes centrosomal MIB1 and HERC2 whereas forcing PCM1 to the centrosome restores their localization, establishing that PCM1 organization dictates centrosomal levels of both ubiquitin ligases. Together, these results demonstrate that pericentrosomal PCM1 accumulation, driven by T21-dependent elevation of PCNT, impairs primary ciliogenesis by sequestering MIB1 and HERC2 from the centrosome.

## RESULTS

### Trisomy 21 delays mother centriole uncapping and primary ciliogenesis

To study the effects of T21 on centriolar uncapping, we used isogenic RPE-1 cell lines engineered with either two (Disomy 21; D21) or three (Trisomy 21; T21) copies of chromosome 21 (Lane et al., 2014). Ciliation and mother centriole uncapping frequency were measured across a serum depletion time course to induce ciliogenesis (Figure 1A). Consistent with prior work, T21 delayed primary ciliogenesis relative to D21 (Galati et al., 2018; Jewett et al., 2023; McCurdy et al., 2022). T21 also delayed centriolar uncapping, reducing uncapping and ciliation frequency at all time points up to 48 hours of serum depletion. By 48 hours, T21 uncapping and ciliation frequencies recovered to D21 levels, indicating a delay rather than permanent impairment. This suggests that the uncapping machinery remains competent in T21 but operates with reduced efficiency, increasing the time required for CP110 removal from the mother centriole. Together, these data indicate that T21-associated ciliogenesis delays arise upstream, at centriolar uncapping.

**Figure 1.**
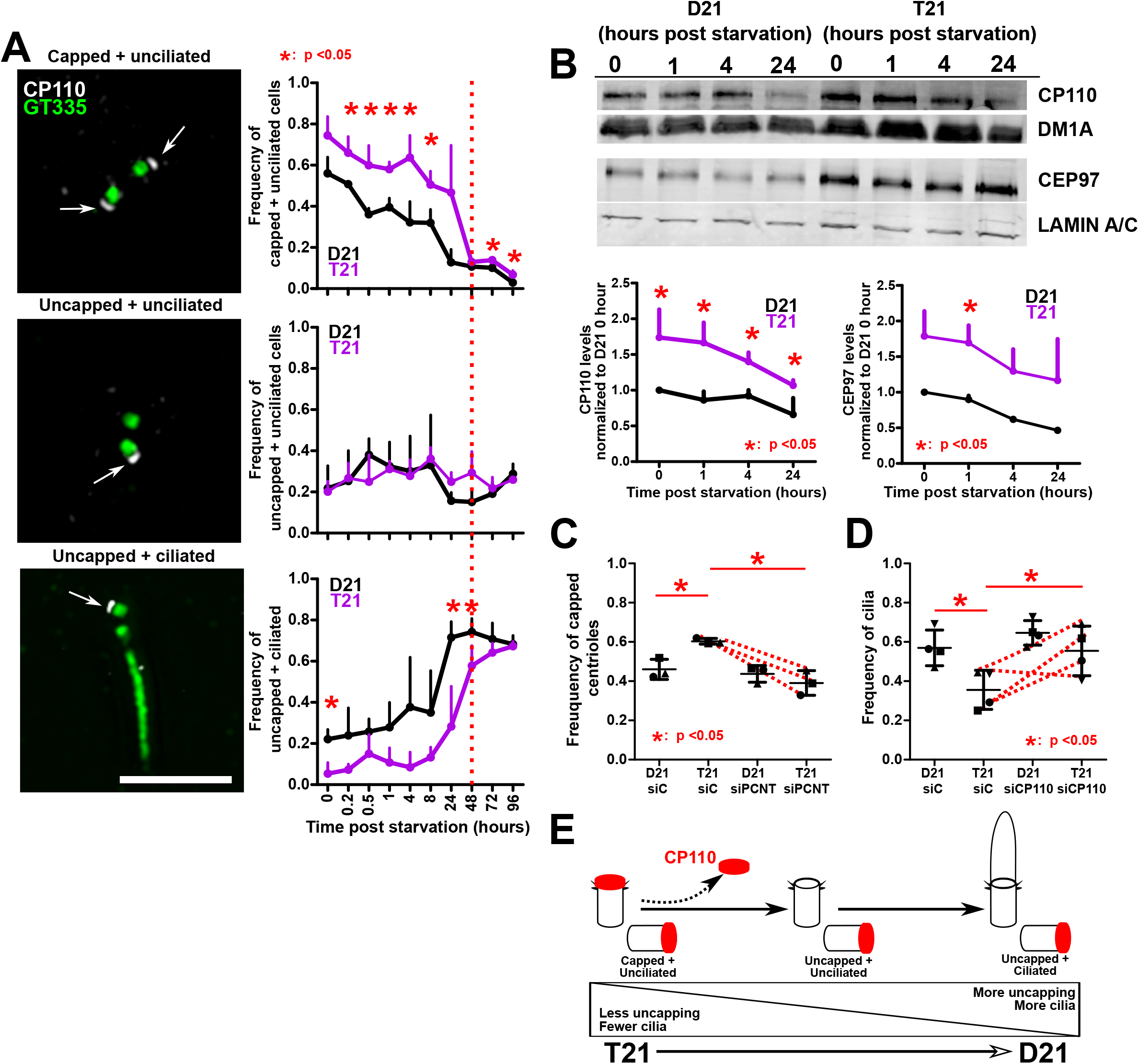
Trisomy 21 delays uncapping and ciliogenesis. (**A**) Left panels, disomy 21 (D21) and trisomy 21 (T21) RPE-1 cells stained for centrioles and cilia (GT335; green) and CP110 (CP110; grayscale). White arrows designate capped centrioles. Scale bar, 5 µm. Right panels, relative frequencies of uncapping and ciliogenesis state across a serum starvation time course in D21 and T21. Mean ± SD. *, p < 0.05. (**B**) T21 increases total levels of CP110 and CEP97 across a serum starvation time course. Top panels, western blots of CP110 and CEP97 with respective loading controls DM1A and Lamin A/C in D21 and T21 across a serum starvation time course. Bottom panels, quantification of CP110 and CEP97 western blot band intensities in D21 and T21. CP110 and CEP97 band intensities were corrected using respective DM1A and Lamin A/C loading controls. Plotted band intensities were normalized to D21 0 hour levels. Mean ± SD. *, p < 0.05. (**C**) T21 perturbs centriolar uncapping in a PCNT dependent manner. Centriolar capping frequencies in D21 and T21 with or without knockdown of PCNT 24 hours after serum starvation. Changes in mean values indicated with red lines. Mean ± SD. *, p < 0.05. (**D**) Reduction of CP110 in T21 rescues the T21 uncapping defect. Centriolar capping frequencies in D21 and T21 with or without knockdown of CP110 24 hours after serum starvation. Changes in mean values indicated with red lines. Mean ± SD. *, p < 0.05. (**E**) T21 increases the proportion of cells that are uncapped and unciliated at ciliogenesis onset.

Centriolar uncapping is triggered by ubiquitination of CP110 and subsequent proteasome-mediated degradation of CP110 and CEP97, which lowers their protein levels upon ciliogenesis induction (Spektor et al., 2007; Xie et al., 2023). To test whether T21 alters this degradation, total levels of CP110 and CEP97 were measured across a serum depletion time course (Figure 1B and Supplemental Figure S1G). Both proteins were elevated at all timepoints post serum depletion in T21 compared to D21. Although both D21 and T21 reduced CP110 (D21:-34.1%, T21:-38.6%) and CEP97 (D21:-53.4%, T21:-34.8%) at 24 hours post serum depletion, protein levels at all timepoints remained higher in T21. In summary, CP110 and CEP97 are elevated and their degradation is perturbed in T21, supporting a model in which reduced CP110 and CEP97 degradation delays centriolar uncapping.

We next asked whether the uncapping defect is PCNT dependent. Reducing PCNT levels to sub-disomic levels in D21 and T21 rescued the uncapping frequency in T21 to D21 levels (Figure 1C and Supplemental Figure S1A), indicating that elevated PCNT delays uncapping and subsequent ciliogenesis in T21. To confirm the uncapping and ciliation delays were not specific to serum depletion, ciliogenesis was induced using KI16245, an LPA receptor inhibitor (Supplemental Figure S1D) (Walia et al., 2019). T21 reduced uncapping and ciliation at all time points after LPA receptor inhibition. PCNT was elevated in T21 across all timepoints with both serum depletion and LPA inhibition (Supplemental Figure S1E and S1F).

Because CP110 levels influence centriolar uncapping and ciliogenesis, we asked whether reducing CP110 rescues the delayed uncapping in T21 (Figure 1D and Supplemental Figure S1B). Reducing CP110 to sub-disomic levels rescued T21 ciliation frequency to D21 levels. To test whether this rescue acts by lowering PCNT levels, we measured centrosomal PCNT levels with or without CP110 knockdown (Supplemental Figure S1C). CP110 reduction did not alter centrosomal PCNT. Thus, directly reducing CP110 rescues ciliogenesis without correcting elevated PCNT, demonstrating that lowering CP110 is sufficient to bypass the T21-associated uncapping defect. Together, these findings support a model in which T21-driven PCNT elevation impairs centriolar uncapping, which in turn delays ciliogenesis (Figure 1E).

### Trisomy 21 reduces centrosomal E3 ligase localization and CP110 ubiquitination

Centriolar uncapping requires CP110 ubiquitination and subsequent proteasome-mediated degradation (Xie et al., 2023). Because CP110 and CEP97 were elevated in T21 (Figure 1B), we asked whether CP110 ubiquitination was altered. We measured total ubiquitinated CP110 4 hours after serum depletion with or without the proteasome inhibitor MG132 using complementary immunoprecipitation approaches (Figure 2A and 2B and Supplemental Figure S2D and S2E). Ubiquitin immunoprecipitation followed by CP110 immunoblotting revealed reduced ubiquitinated CP110 in T21 following MG132 treatment (Figure 2A and Supplemental Figure S2D). Reciprocal CP110 immunoprecipitation showed a similar overall trend, although changes were less apparent and not statistically significant. This may reflect a broader population of ubiquitinated proteins and CP110-associated complexes recovered by CP110 immunoprecipitation, which could obscure changes in ubiquitinated CP110 (Figure 2B and Supplemental Figure S2E). Together, these data support a model in which impaired CP110 ubiquitination reduces CP110 and CEP97 degradation at the mother centriole, delaying centriolar uncapping and ciliogenesis.

**Figure 2.**
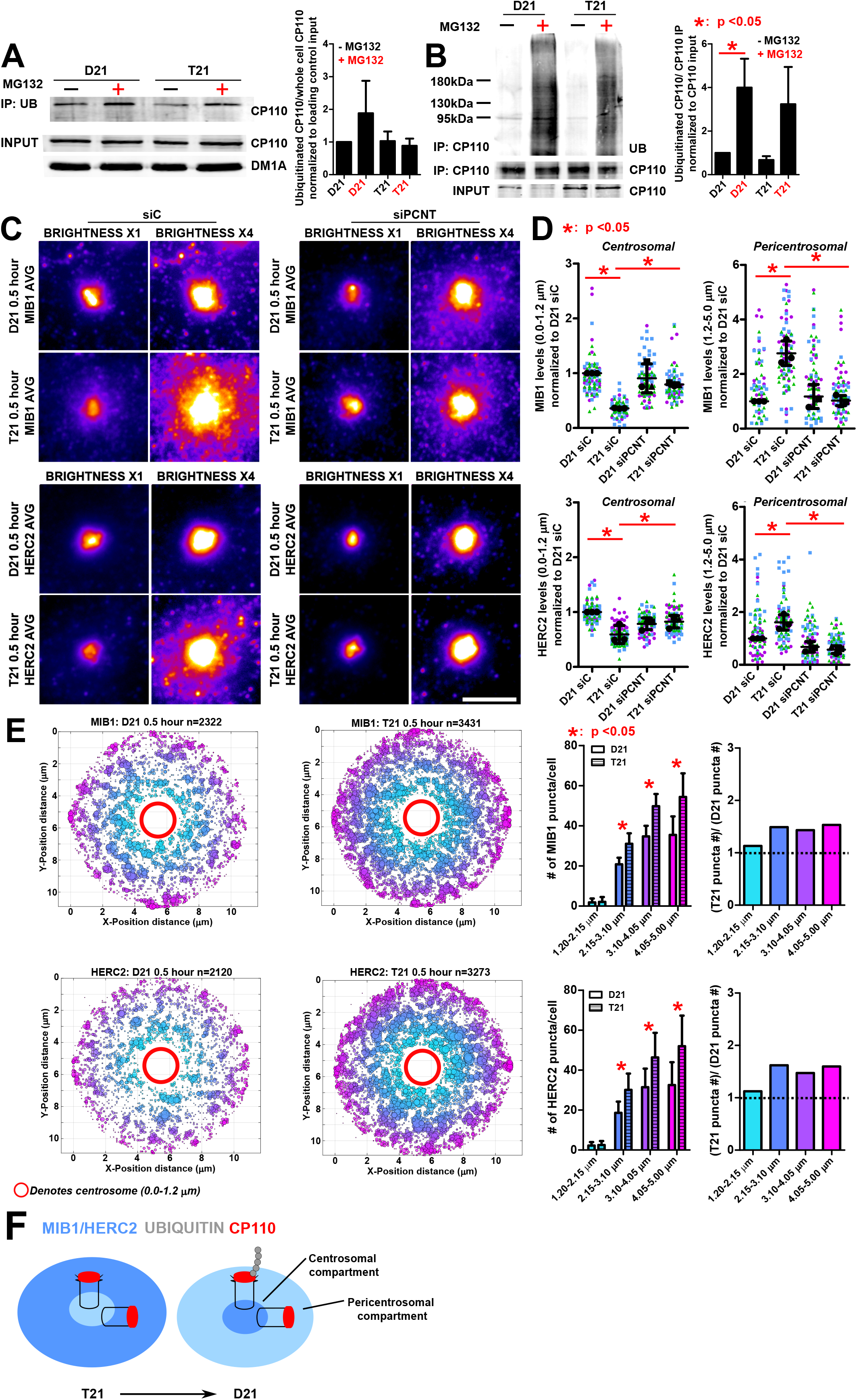
Trisomy 21 reduces CP110 ubiquitination and centrosomal localization of E3 ligases. (**A**) Left panel, western blot of CP110 in D21 and T21 after immunoprecipitation (IP) of ubiquitin with or without the presence of proteasomal inhibitor MG132. CP110 and DM1A inputs displayed below IP blot. Cells were starved for 4 hours before harvesting lysate. Right panel, quantification of ubiquitinated CP110 normalized to CP110 and loading control input bands. All band values were normalized to D21 (-MG132). Mean ± SD. *, p < 0.05. (**B**) Left panel, western blot of ubiquitin and CP110 in D21 and T21 after immunoprecipitation (IP) of CP110 with or without the presence of proteasomal inhibitor MG132. CP110 inputs displayed below IP blot. Cells were starved for 4 hours before harvesting lysate. Right panel, quantification of ubiquitinated CP110 normalized to CP110 IP and input bands. All band values were normalized to D21 (-MG132). Mean ± SD. *, p < 0.05. (**C**) T21 reduces centrosomal levels of MIB1 and HERC2 and increases pericentrosomal levels of MIB1 and HERC2 in a PCNT dependent manner. Image averages of D21 and T21 cells stained for MIB1 or HERC2 with or without knockdown of PCNT 0.5 hours after serum starvation. Scale bar, 5 µm. (**D**) Quantification of centrosomal and pericentrosomal MIB1 and HERC2 in D21 and T21 with or without knockdown of PCNT 0.5 hours after serum starvation. MIB1 and HERC2 levels were normalized to D21 siC averages. Mean ± SD. *, p < 0.05. (**E**) T21 increases pericentrosomal MIB1 and HERC2 puncta number 0.5 hours after serum starvation. Left panels, pericentrosomal MIB1 and HERC2 puncta colored as a function of distance from the centrosome. Size of puncta correlates with puncta density in that region. Right panels, quantification of puncta number in D21 and T21. Mean ± SD. *, p < 0.05. (**F**) T21 shifts MIB1 and HERC2 partitioning away from the centrosome, reducing centrosomal E3 ligase availability and CP110 ubiquitination relative to D21.

T21-dependent PCNT elevation promotes the accumulation of pericentrosomal assemblies that mislocalize ciliogenesis regulators (Galati et al., 2018; Jewett et al., 2023; McCurdy et al., 2022). One model is that PCNT assemblies bind and sequester ciliogenesis proteins away from the centrosome. Because T21 reduces CP110 ubiquitination, we asked whether T21 and elevated PCNT mislocalizes the CP110 E3 ubiquitin ligases MIB1 and HERC2. We measured centrosomal and pericentrosomal MIB1 and HERC2 in D21 and T21 0.5 hours after serum depletion (Figure 2C and 2D). T21 reduced centrosomal MIB1 and HERC2 while increasing both proteins in pericentrosomal regions, where they manifested as puncta surrounding the centrosome (Figure 2C-2E). Enrichment varied across pericentrosomal regions, but nearly all regions in T21 showed more puncta than D21, consistent with T21 PCNT assemblies sequestering MIB1 and HERC2 (Figure 2E).

To test whether reduced centrosomal MIB1 and HERC2 in T21 reflect lower total cellular levels, we measured whole cell MIB1 and HERC2 fluorescence (Supplemental Figure S2A and S2B). Both proteins were elevated in T21, arguing against reduced total levels as the cause of their centrosomal depletion. To test whether mislocalization of another CP110 uncapping factor contributes to the T21 phenotype, we measured centrosomal TTBK2, a kinase essential for uncapping (Supplemental Figure S2C) (Goetz et al., 2012). Centrosomal TTBK2 was unchanged in T21 compared to D21, indicating that the disruption of the CP110 uncapping machinery is not a general consequence of altered centrosomal protein localization and instead points to selective mislocalization of MIB1 and HERC2.

To test whether the centrosomal loss and pericentrosomal accumulation of MIB1 and HERC2 are PCNT dependent, we reduced PCNT to sub-disomic levels in D21 and T21 (Figure 2C and 2D). In T21, reduced PCNT rescued both centrosomal MIB1 and HERC2 and their elevated pericentrosomal accumulation to D21 levels. Collectively, these results support a model in which pericentrosomal PCNT assemblies sequester MIB1 and HERC2, depleting them from the centrosome and impairing CP110 ubiquitination and uncapping (Figure 2F).

### Trisomy 21 increases PCNT and PCM1 pericentrosomal accumulation during centriolar uncapping

Like MIB1 and HERC2, elevated PCNT increases pericentrosomal PCM1 (Jewett et al., 2023; McCurdy et al., 2022). To test whether PCM1 accumulates at PCNT foci during uncapping, we measured PCM1-PCNT colocalization during the serum depletion time course (Figure 3A). PCNT and PCM1 colocalized more frequently at both centrosomal and pericentrosomal regions in T21 than D21. PCM1 was also elevated in T21 at all time points of serum depletion and LPA receptor inhibition (Supplemental Figure S3B), and T21 increased PCNT and PCM1 puncta number at nearly every pericentrosomal region 30 minutes post serum starvation (Figure 3C). Reducing PCNT to sub-disomic levels reduced pericentrosomal PCM1 accumulation, confirming that it is dependent on elevated PCNT (Figure 3B). Consistent with prior work, PCM1 reduction impaired ciliogenesis in D21 (Supplemental Figure S3D) (Hall et al., 2023). Surprisingly, PCM1 reduction in T21 increased ciliogenesis. Although the mechanism remains unknown, PCM1 reduction in T21 reduced centrosomal PCNT levels to D21 levels, raising the possibility that PCM1 binding stabilizes PCNT and promotes its accumulation (Supplemental Figure S3D).

**Figure 3.**
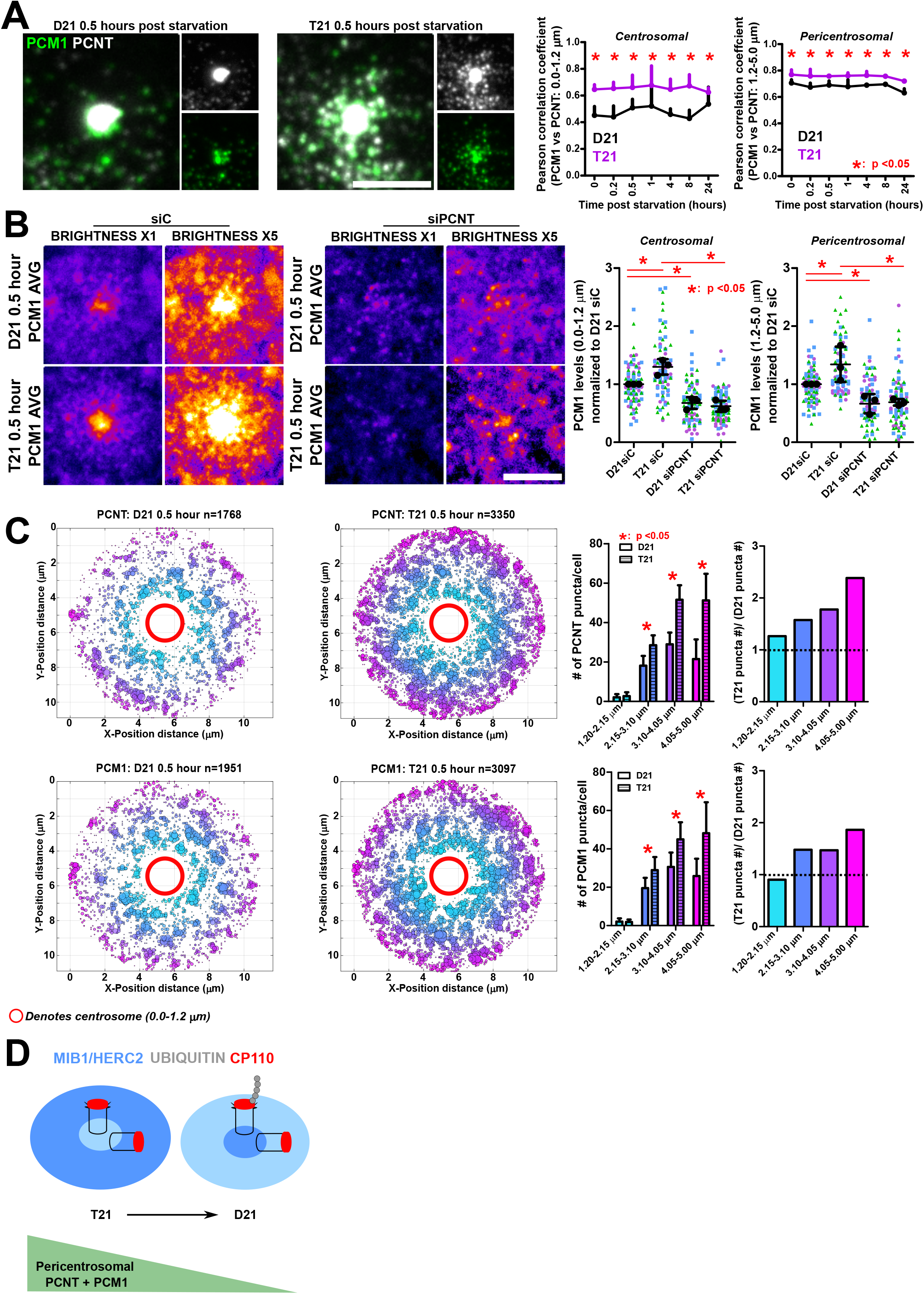
Trisomy 21 increases PCNT and PCM1 pericentrosomal accumulation during centriolar uncapping. (**A**) T21 increases PCM1 and PCNT colocalization at centrosomal and pericentrosomal regions. Left panels, D21 and T21 cell stained for PCNT (grayscale) and PCM1 (green) 0.5 hours post serum starvation. Scale bar, 5 µm. Right panels, Pearson correlation coefficients for PCM1 and PCNT in D21 and T21 in centrosomal and pericentrosomal regions across a serum starvation time course. Mean ± SD. *, p < 0.05. (**B**) Centrosomal and pericentrosomal accumulation of PCM1 in T21 is PCNT dependent. Left panels, image averages of D21 and T21 cells stained for PCM1 with or without knockdown of PCNT 0.5 hours post serum starvation. Scale bar, 5 µm. Right panel, quantification of centrosomal and pericentrosomal PCM1 levels with or without knockdown of PCNT 0.5 hours post serum starvation. Levels were normalized to D21 siC averages. Mean ± SD. *, p < 0.05. (**C**) T21 increases pericentrosomal PCNT and PCM1 puncta number 0.5 hours after serum starvation. Left panels, pericentrosomal PCNT and PCM1 puncta colored as a function of distance from the centrosome. Size of puncta correlates with puncta density in that region. Right panels, quantification of puncta number in D21 and T21. Mean ± SD. *, p < 0.05. (**D**) Elevated PCNT in T21 increases pericentrosomal PCM1 accumulation, redistributing MIB1 and HERC2 away from the centrosome and reducing CP110 ubiquitination.

To test whether pericentrosomal PCM1 drives pericentrosomal MIB1 and HERC2 accumulation, we quantified PCM1 colocalization with each E3 ligase (Supplemental Figure S3A). PCM1 colocalized with both ligases, and more strongly with MIB1. Average colocalization was equivalent in D21 and T21, indicating that increased PCM1 accumulation does not change the frequency of PCM1 association with MIB1 or HERC2 (Supplemental Figure S3A).

Because reducing CP110 rescues uncapping and ciliogenesis in T21, we tested whether CP110 reduction modulates PCM1. Reducing CP110 decreased centrosomal and pericentrosomal PCM1 in both D21 and T21 (Supplemental Figure S3C). This suggests that CP110 reduction rescues T21 uncapping through two mechanisms: lowering PCM1 and bypassing the uncapping requirement. Together, these data support a model in which T21-driven PCNT elevation expands pericentrosomal PCM1 puncta in the same regions where MIB1 and HERC2 are mislocalized (Figure 3D).

### T21 does not alter PCNT or PCM1 motility to the centrosome

Both PCNT and PCM1 interact with MT-dependent transport machinery and are thought to move bound cargoes to and from the centrosome along MTs (Dammermann & Merdes, 2002; Galati et al., 2018; Kubo et al., 1999; Lee & Rhee, 2011; McCurdy et al., 2022; Purohit et al., 1999; Tollenaere et al., 2015). Recent studies have refined this model, revealing heterogeneous centriolar satellite dynamics in which directed, MT-dependent movement occurs alongside more frequent, non-directional, diffusive movement (Begar et al., 2025; Conkar et al., 2019; Pachinger et al., 2025). Detailed characterization of PCM1 motility showed that most PCM1-positive centriolar satellites display non-directional, diffusive movement (Conkar et al., 2019). In T21, one possibility is that elevated pericentrosomal PCNT impairs transport of PCM1, and its MIB1 and HERC2 cargoes, to the centrosome. Consistent with this, cells with elevated pericentrosomal PCM1 had less centrosomal MIB1 and HERC2 (Figure 4A). To test whether T21 alters PCNT or PCM1 transport, we visualized PCNT (endogenous PCNT-mNeon) and PCM1 (exogenous PCM1-GFP) in live cells. Because overexpression of PCM1 increases centrosomal and pericentrosomal PCNT in D21 cells (Figure 5C), a doxycycline titration curve identified 18 nM as a PCM1-GFP concentration that does not affect PCNT levels (Supplemental Figure S4A). Both PCNT-mNeon and PCM1-GFP localized to centrosomal and pericentrosomal regions, as expected (Figure 4B and Supplemental Figure S4B).

**Figure 4.**
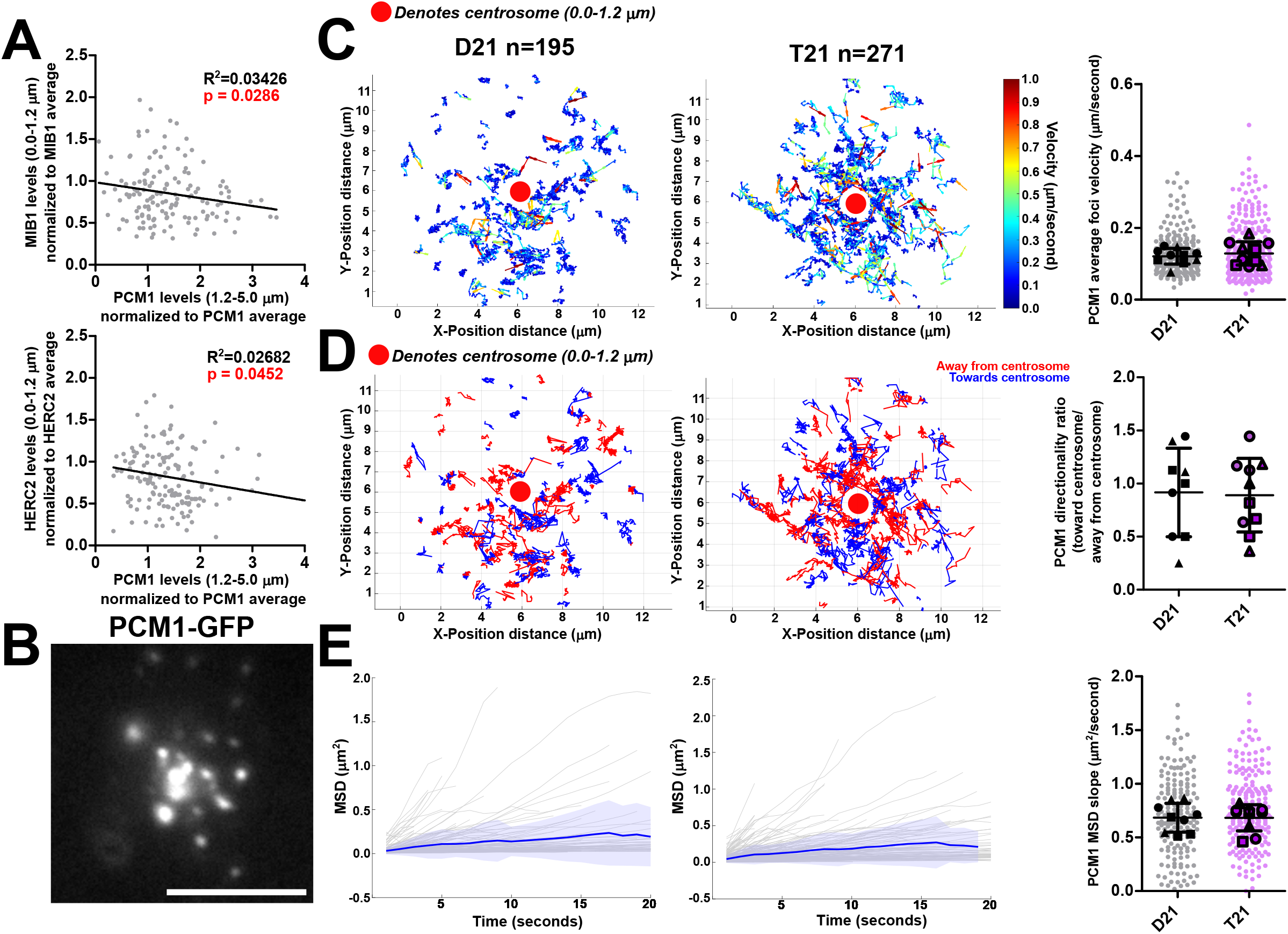
Trisomy 21 does not alter PCNT or PCM1 motility. (**A**) Pericentrosomal PCM1 levels negatively correlate with centrosomal MIB1 and HERC2 levels. D21 and T21 values were combined on the same plot. p-values denoted in red. R^2^ values denoted in black. (**B**) Inducible overexpression of PCM1 targets PCM1-GFP to the centrosome and pericentrosomal regions. Scale bar, 5 µm. (**C**) T21 does not alter average PCM1 puncta velocity. Left panels, plots showing average PCM1 track trajectories in the pericentrosomal region 0.5 hours post serum starvation. Trajectories are colored by average speed (µm/sec). Right panel, average PCM1 foci velocities in D21 and T21. Mean ± SD. *, p < 0.05. (**D**) T21 does not alter PCM1 puncta trajectory direction. Left panels, plots showing average PCM1 track trajectories in the pericentrosomal region 0.5 hours post serum starvation. Trajectories are colored by direction of movement either toward or away from the centrosome. Right panel, PCM1 foci directionality ratios in D21 and T21. Mean ± SD. *, p < 0.05. (**E**) T21 does not alter PCM1 mean squared displacement (MSD). Left panels, individual MSD trajectories for PCM1 puncta in D21 and T21 0.5 hours post serum starvation. Right panel, PCM1 MSD slopes for D21 and T21. Mean ± SD. *, p < 0.05.

**Figure 5.**
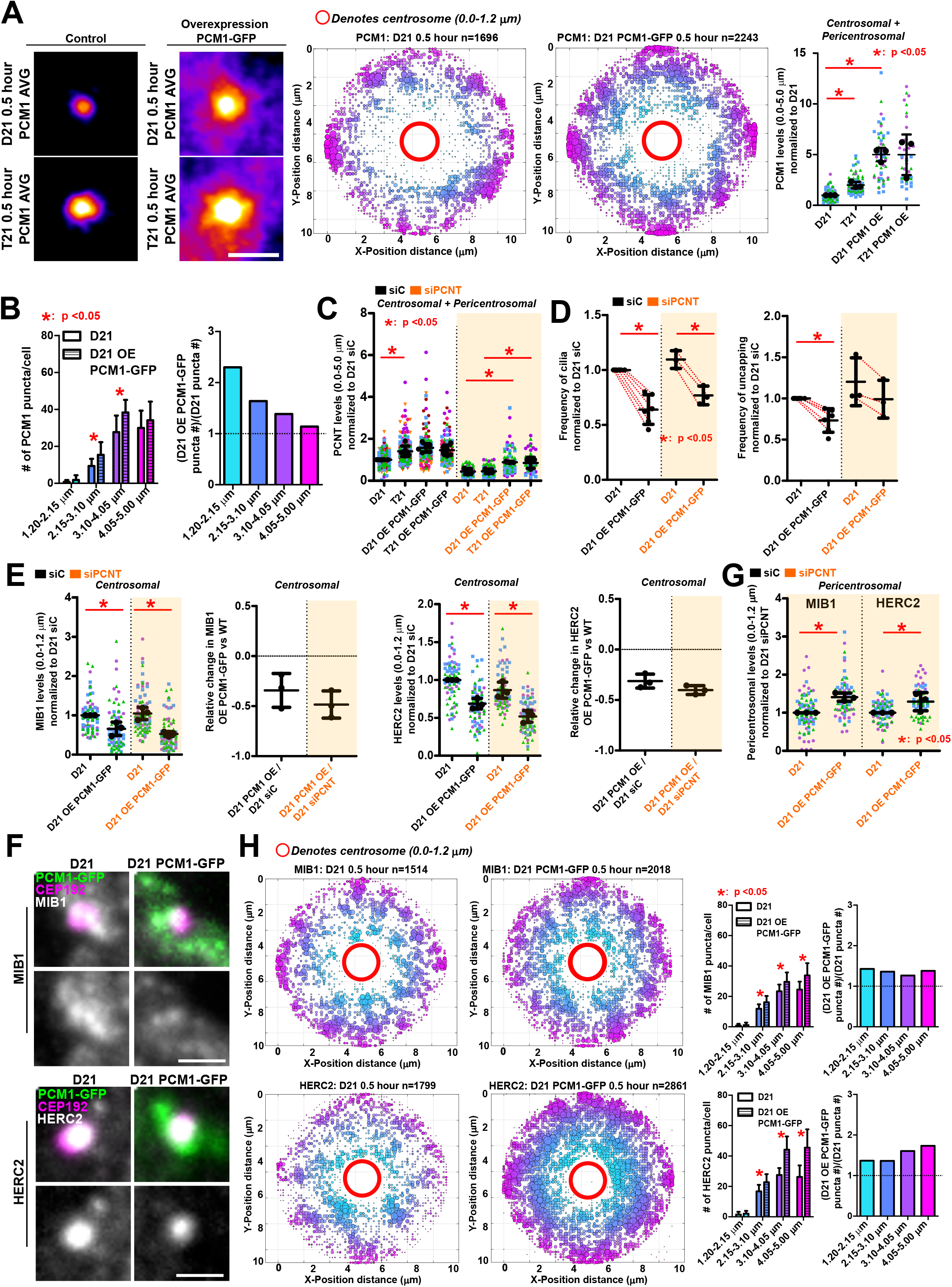
Elevated PCM1 mislocalizes MIB1 and HERC2 and causes uncapping and ciliogenesis defects. (**A**) PCM1 overexpression increases centrosomal and pericentrosomal levels of PCM1. Left panels, image averages of D21 and T21 cells stained for PCM1 with or without overexpression of PCM1 0.5 hours after serum starvation. Scale bar, 5 µm. Middle panels, pericentrosomal PCM1 puncta colored as a function of distance from the centrosome in D21 cells with or without overexpression of PCM1. Size of puncta correlates with puncta density in that region. Right panel, quantification of centrosomal and pericentrosomal PCM1 levels in D21 and T21 with or without PCM1 overexpression. Levels were normalized to D21 non-overexpressing cells. Mean ± SD. *, p < 0.05. (**B**) PCM1 overexpression increases PCM1 pericentrosomal puncta number. Quantification of PCM1 puncta number in D21 cells with or without PCM1 overexpression 0.5 hours post serum starvation. Mean ± SD. *, p < 0.05. (**C**) PCM1 overexpression increases centrosomal and pericentrosomal PCNT. Quantification of centrosomal and pericentrosomal PCNT in D21 and T21 cells 0.5 hours post serum starvation with or without knockdown of PCNT. Levels were normalized to D21 siC averages. Mean ± SD. *, p < 0.05. (**D**) PCM1 overexpression reduces centriolar uncapping and ciliogenesis in D21 with or without knockdown of PCNT. Left panel, quantification of cilia frequency 24 hours post serum starvation in D21 cells with or without knockdown of PCNT. Frequencies were normalized to D21 siC averages. Mean ± SD. *, p < 0.05. Right panel, quantification of uncapping frequency 24 hours post serum starvation in D21 cells with or without knockdown of PCNT. Frequencies were normalized to D21 siC averages. Mean ± SD. *, p < 0.05. (**E**) PCM1 overexpression reduces centrosomal MIB1 and HERC2 levels in D21 0.5 hours post serum starvation with or without knockdown of PCNT. Left panels, quantification of centrosomal MIB1 levels and relative change in MIB1 0.5 hours post serum starvation in D21 cells with or without knockdown of PCNT. Levels were normalized to D21 siC averages. Mean ± SD. *, p < 0.05. Right panels, quantification of centrosomal HERC2 levels and relative change in HERC2 0.5 hours post serum starvation in D21 cells with or without knockdown of PCNT. Levels were normalized to D21 siC averages. Mean ± SD. *, p < 0.05. (**F**) PCM1 overexpression reduces centrosomal MIB1 and HERC2 levels in D21 0.5 hours post serum starvation with knockdown of PCNT. Top panels, D21 cells stained for CEP192 (magenta) and MIB1 (grayscale) with or without overexpression of PCM1 (PCM1-GFP) 0.5 hours after serum starvation with knockdown of PCNT. Scale bar, 2.5 µm. Bottom panels, D21 cells stained for CEP192 (magenta) and HERC2 (grayscale) with or without overexpression of PCM1 (PCM1-GFP) 0.5 hours after serum starvation with knockdown of PCNT. Scale bar, 2.5 µm. (**G**) PCM1 overexpression increases pericentrosomal MIB1 and HERC2 levels in D21 0.5 hours post serum starvation with knockdown of PCNT. Quantification of pericentrosomal MIB1 and HERC2 levels 0.5 hours post serum starvation in D21 cells with knockdown of PCNT. Levels were normalized to D21 siC averages. Mean ± SD. *, p < 0.05. (**H**) PCM1 overexpression increases MIB1 and HERC2 pericentrosomal puncta in D21 0.5 hours post serum starvation. Left panels, pericentrosomal MIB1 and HERC2 puncta colored as a function of distance from the centrosome. Size of puncta correlates with puncta density in that region. Right panels, quantification of MIB1 and HERC2 puncta number in D21 and T21. Mean ± SD. *, p < 0.05.

To test whether T21 alters PCNT or PCM1 transport to the centrosome, we measured average pericentrosomal puncta velocities in D21 and T21 cells 0.5 hours post serum depletion (Figure 4C and Supplemental Figure S4C-S4D). T21 PCNT foci showed a small but significant velocity reduction, whereas PCM1 foci velocities were unchanged. The frequency of PCNT and PCM1 puncta movement toward or from the centrosome was also similar in D21 and T21, indicating no directional bias away from the centrosome in T21 (Figure 4D and Supplemental Figure S4C-S4D).

To assess whether fewer puncta were actively moving, we measured mean squared displacement (MSD) (Figure 4E and Supplemental Figure S4C-S4D). Most PCNT and PCM1 puncta exhibited constrained diffusive transport (MSD slope <1.0), indicating that the majority of puncta move with constrained diffusion rather than actively transporting to or from the centrosome, consistent with previous observations (Conkar et al., 2019). Finally, fluorescence recovery after photobleaching (FRAP) of PCNT and PCM1 revealed no differences in the mobile fraction or recovery kinetics between D21 and T21 cells (Supplemental Figure S4E), indicating that elevated PCNT does not alter the dynamic exchange of PCM1 within the pericentrosomal compartment despite its increased accumulation. In D21 and T21, PCM1 fluorescence recovered to approximately 50% of prebleach levels, indicating that PCM1 exists as both a dynamically exchanging pool and as a stable or slowly exchanging pool. The relative contributions of these populations were unchanged in T21, suggesting that elevated PCNT does not alter the balance between dynamic and stable PCM1 populations. Although FRAP cannot distinguish whether the dynamic or stable PCM1 pool mediates the pericentrosomal capture of MIB1 and HERC2, these findings argue against a model in which elevated PCNT promotes PCM1 accumulation by altering transport or stabilizing PCM1 interactions. Instead, the increased abundance of PCM1 is consistent with expansion of the pericentrosomal compartment while preserving the dynamic behavior of its constituent PCM1 populations. Collectively, these results suggest that, during ciliogenesis, most pericentrosomal PCNT and PCM1 puncta do not actively transport cargoes to the centrosome, and that elevated PCNT and PCM1 do not alter PCM1 trafficking. Furthermore, in the context of T21, this argues that reduced centrosomal MIB1 and HERC2 arises from sequestration by increased pericentrosomal PCM1.

### Elevated PCM1 mislocalizes MIB1 and HERC2 and causes uncapping and ciliogenesis defects

T21-driven PCNT elevation promotes the accumulation of PCM1 in the pericentrosomal region, creating additional sites for association with its binding partners MIB1 and HERC2. This redistribution of PCM1 is accompanied by the mislocalization of MIB1 and HERC2 away from the centrosome. To test whether increased PCM1 alone is sufficient to mislocalize centrosomal MIB1 and HERC2, we overexpressed PCM1 in D21 and T21 cells (Figure 5A). PCM1 overexpression produced an approximately 5-fold increase in PCM1 protein at and peripheral to the centrosome, and increased pericentrosomal PCM1 puncta number (Figure 5A-5B). Because PCNT reduction lowers PCM1, we also tested how PCM1 overexpression affects PCNT (Figure 5C). PCM1 overexpression raised centrosomal and pericentrosomal PCNT to T21 levels in D21 but did not further raise PCNT levels in T21, suggesting that PCM1 promotes PCNT accumulation up to a limit that T21 has already reached (Figure 5C). To determine whether elevated PCM1 is sufficient to perturb uncapping and ciliogenesis independently of elevated PCNT, we overexpressed PCM1 following PCNT reduction. Notably, PCM1 overexpression maintained PCNT levels at D21 siControl levels in both D21 and T21 cells (Figure 5C), suggesting that increased PCM1 may promote PCNT accumulation or stability. Importantly, PCNT reduction did not prevent the formation of pericentrosomal PCM1 assemblies in PCM1 overexpression contexts (Figure S5E). PCM1 overexpression reduced centriolar uncapping and ciliation frequency in D21 under both normal and reduced PCNT conditions (Figure 5D and Supplemental Figure S5A), demonstrating that elevated PCM1 is sufficient to reduce uncapping. PCM1 overexpression in D21 also reduced centrosomal MIB1 and HERC2 while increasing pericentrosomal levels and number of puncta in normal and reduced PCNT conditions (Figure 5E-5H). Similar effects of PCM1 overexpression were observed in T21 cells, including reduced ciliation and centrosomal MIB1 and HERC2 (Supplemental Figure S5B-S5D). Together, these findings support a model in which elevated PCM1 is sufficient to recapitulate key consequences of T21-associated PCNT elevation, linking increased PCNT to MIB1 and HERC2 redistribution and subsequent defects in centriolar uncapping and ciliogenesis.

### Repositioning PCM1 relocalizes MIB1 and HERC2

To test whether MIB1 and HERC2 localization follows PCM1, we first altered PCM1 localization pharmacologically (Figure S5F). Treatment with the proteasome inhibitor MG132 induces aggresome formation, a cellular response that causes accumulation of proteins at the centrosome. We therefore used MG132-induced aggresome formation to increase PCM1 localization in the centrosomal region and asked whether MIB1 and HERC2 were similarly redistributed. MG132 treatment increased centrosomal PCM1, MIB1, and HERC2 in D21, with more variable responses observed in T21 (Supplemental Figure S5F). Conversely, depolymerizing MTs with nocodazole to disperse PCM1 reduced centrosomal PCM1, MIB1, and HERC2 in D21. In T21, nocodazole reduced centrosomal PCM1 and MIB1, but not HERC2. Together, these data suggest that altering the spatial distribution of PCM1 is sufficient to redistribute centrosomal MIB1 and HERC2.

To directly test whether PCM1 localization determines the centrosomal localization of MIB1 and HERC2, we adapted an Anchor-Away system (Aydin Ö et al., 2020; Edwards & Wandless, 2007) to direct PCM1 either to or away from the centrosome. Stable D21 and T21 cell lines were engineered to co-express PCM1-GFP-FKBP and the dynein adaptor BICD2-FRB. Rapamycin promotes the interaction of FKBP and FRB (Edwards & Wandless, 2007), tethering PCM1-GFP-FKBP to BICD2-FRB. Because BICD2 engages the dynein transport machinery, this induced interaction drives PCM1 toward the microtubule minus ends at the centrosome, resulting in its centrosomal enrichment. Rapamycin treatment accordingly increased centrosomal PCM1 in both D21 and T21 cells (Figure 6A and 6C). Centrosomal MIB1 and HERC2 also increased in both cell lines, indicating that increasing PCM1 at the centrosome is sufficient to increase the centrosomal localization of these ligases. Importantly, increased centrosomal PCM1, MIB1, and HERC2 increased ciliation frequencies in D21 and T21 (Figure 6C), linking centrosomal localization of PCM1-associated ligases with ciliogenesis.

**Figure 6.**
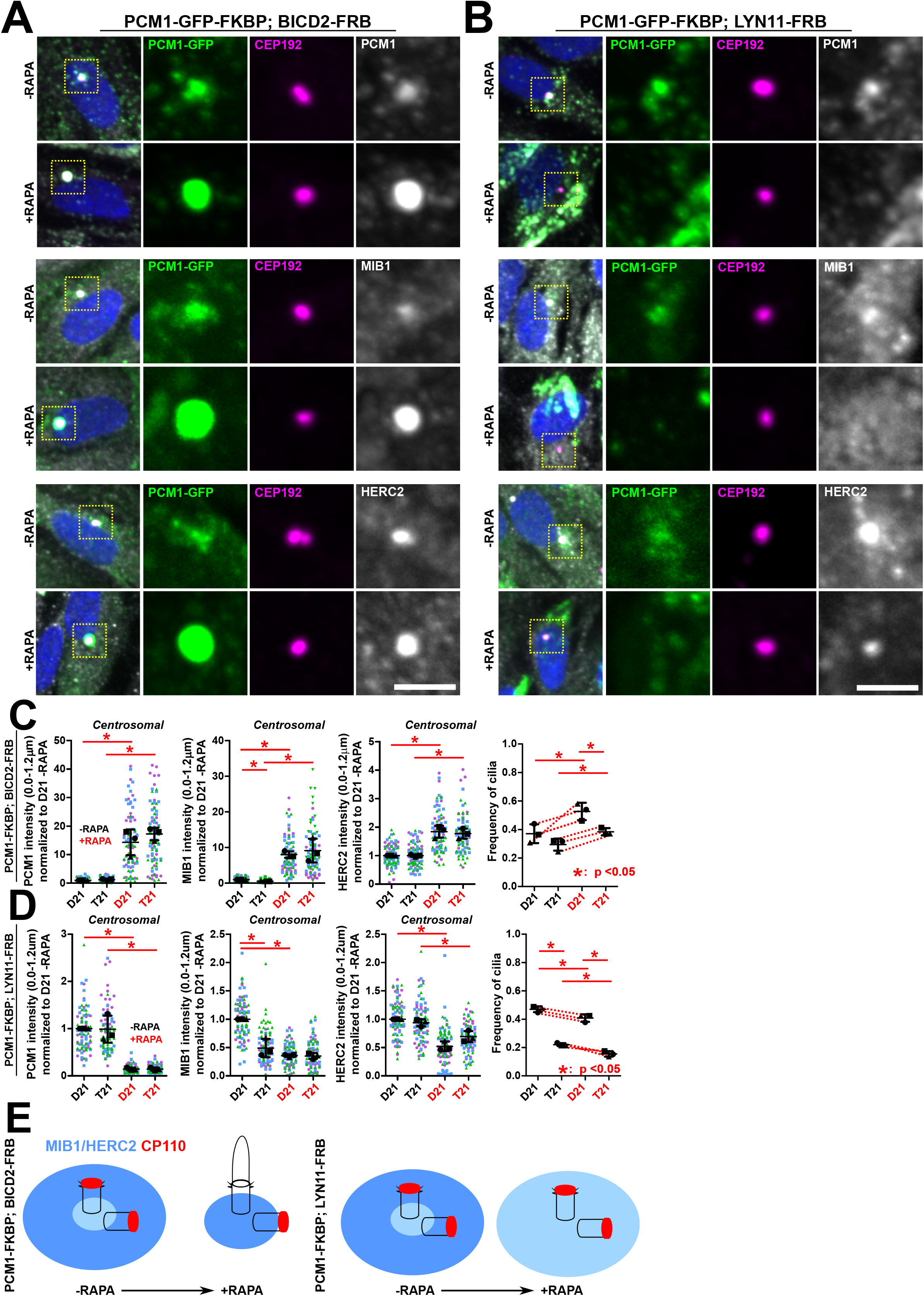
Repositioning PCM1 relocalizes MIB1 and HERC2. (**A**) Pulling PCM1 to the centrosome increases centrosomal PCM1, MIB1, and HERC2 levels. Top panel, D21 cell expressing PCM1-GFP-FKBP; BICD2-FRB and stained for PCM1 (PCM1; grayscale) and CEP192 (magenta) with or without the presence of rapamycin. Middle panel, D21 cell expressing PCM1-GFP-FKBP; BICD2-FRB stained for MIB1 (grayscale) and CEP192 (magenta) with or without the presence of rapamycin. Bottom panel, D21 cell expressing PCM1-GFP-FKBP; BICD2-FRB stained for HERC2 (grayscale) and CEP192 (magenta) with or without the presence of rapamycin. Scale bar, 5 µm. (**B**) Pulling PCM1 away from the centrosome decreases centrosomal PCM1, MIB1, and HERC2 levels. Top panel, D21 cell expressing PCM1-GFP-FKBP; LYN11-FRB stained for PCM1 (PCM1 AB; grayscale) and CEP192 (magenta) with or without the presence of rapamycin. Middle panel, D21 cell expressing PCM1-GFP-FKBP; LYN11-FRB stained for MIB1 (grayscale) and CEP192 (magenta) with or without the presence of rapamycin. Bottom panel, D21 cell expressing PCM1-GFP-FKBP; LYN11-FRB stained for HERC2 (grayscale) and CEP192 (magenta) with or without the presence of rapamycin. Scale bar, 5 µm. (**C**) Pulling PCM1 to the centrosome increases centrosomal PCM1, MIB1, and HERC2 levels and increases cilia frequency. Left panel, centrosomal PCM1 levels 24 hours post serum starvation in D21 and T21 cells expressing PCM1-GFP-FKBP; BICD2-FRB with or without the presence of rapamycin. Middle left panel, centrosomal MIB1 levels 24 hours after serum starvation in D21 and T21 cells expressing PCM1-GFP-FKBP; BICD2-FRB with or without the presence of rapamycin. Middle right panel, centrosomal HERC2 levels 24 hours after serum starvation in D21 and T21 cells expressing PCM1-GFP-FKBP; BICD2-FRB with or without the presence of rapamycin. All values were normalized to D21-rapamycin averages. Right panel, cilia frequencies in D21 and T21 cells expressing PCM1-GFP-FKBP; BICD2-FRB with or without the presence of rapamycin. Changes in mean values indicated with red lines. Mean ± SD. *, p < 0.05. (**D**) Pulling PCM1 away from the centrosome decreases centrosomal PCM1, MIB1, and HERC2 levels and decreases cilia frequency. Left panel, centrosomal PCM1 levels 24 hours post serum starvation in D21 and T21 cells expressing PCM1-GFP-FKBP; LYN11-FRB with or without the presence of rapamycin. Middle left panel, centrosomal MIB1 levels 24 hours after serum starvation in D21 and T21 cells expressing PCM1-GFP-FKBP; LYN11-FRB with or without the presence of rapamycin. Middle right panel, centrosomal HERC2 levels 24 hours after serum starvation in D21 and T21 cells expressing PCM1-GFP-FKBP; LYN11-FRB with or without the presence of rapamycin. All values were normalized to D21-rapamycin averages. Right panel, cilia frequencies in D21 and T21 cells expressing PCM1-GFP-FKBP; LYN11-FRB with or without the presence of rapamycin. Changes in mean values indicated with red lines. Mean ± SD. *, p < 0.05. (**E**) Forced redistribution of PCM1 toward or away from the centrosome correspondingly alters centrosomal MIB1 and HERC2 partitioning and promotes or inhibits ciliogenesis.

To test the effects of directing PCM1 away from the centrosomal and pericentrosomal compartments, stable D21 and T21 cell lines co-expressing PCM1-GFP-FKBP and the plasma membrane-targeted LYN11-FRB (Lindner et al., 2011) were engineered. Rapamycin-induced FKBP–FRB heterodimerization recruits PCM1 to the plasma membrane, depleting it from the centrosomal and pericentrosomal compartments. Consistent with this redistribution, rapamycin treatment reduced centrosomal PCM1 in both D21 and T21 cells (Figure 6B and 6D). Centrosomal MIB1 and HERC2 were also reduced in D21. In T21, centrosomal HERC2 was reduced, whereas MIB1 was unchanged, potentially reflecting its already low basal centrosomal levels observed by overexpression of PCM1. Reducing PCM1 at the centrosome also reduced ciliation frequency in D21 and T21 (Figure 6D). Thus, increasing or decreasing PCM1 proximity to the centrosome produces a corresponding increase or decrease in centrosomal MIB1 and HERC2 and ciliogenesis. Notably, the reduced magnitude of HERC2 responses relative to MIB1 may reflect its weaker association with PCM1, consistent with the lower PCM1-HERC2 colocalization observed (Supplemental Figure S3A). Rapamycin treatment alone did not alter PCNT levels or ciliation frequency (Figure S5G), excluding rapamycin itself as the cause of these effects. Moreover, redirecting PCM1 either toward or away from the centrosome did not alter centrosomal PCNT levels (Figure S5H), indicating that PCM1 redistribution modulates centrosomal MIB1 and HERC2 localization without altering the underlying PCNT scaffold.

Collectively, these results demonstrate that the spatial distribution of PCM1 determines the centrosomal localization of MIB1 and HERC2. Our findings support a model in which PCM1 assemblies spatially partition these ligases by binding and sequestering them, thereby limiting their localization at the centrosome. When PCM1 is redistributed to the centrosome, MIB1 and HERC2 are similarly redistributed; conversely, redistribution of PCM1 away from the centrosome reduces MIB1 and HERC2 localization at the centrosome. In T21 cells, elevated PCNT promotes the expansion of the pericentrosomal PCM1 compartment, increasing sequestration of MIB1 and HERC2 away from the centrosome and altering the local protein environment required for normal ciliogenesis (Figure 6E).

## DISCUSSION

In this study, we identify delayed centriolar uncapping as a previously unrecognized mechanism underlying the ciliogenesis defects in T21. We show that elevated PCNT drives pericentrosomal accumulation of PCM1, which depletes E3 ubiquitin ligases MIB1 and HERC2 from the centrosome, reduces CP110 ubiquitination, and delays its removal from the mother centriole. Reducing PCNT restores centrosomal MIB1 and HERC2, CP110 uncapping, and ciliogenesis, whereas PCM1 overexpression phenocopies each defect independently of PCNT. This establishes pericentrosomal PCM1 accumulation as the critical mediator of these effects. Together, these findings define spatial partitioning as a mechanism by which altered centrosomal and pericentrosomal organization in T21 sequesters the CP110 ubiquitination machinery away from the mother centriole, linking pericentrosomal architecture to the control of ciliogenesis.

Our previous studies proposed that elevated PCNT disrupts primary ciliogenesis by expanding the PCM, increasing centrosomal and acentrosomal microtubule nucleation, and perturbing microtubule-dependent trafficking of ciliary proteins (Galati et al., 2018; Jewett et al., 2023; McCurdy et al., 2022). Our live-cell imaging now argues against altered PCM1 transport as the primary mechanism underlying its accumulation in T21. PCNT and PCM1 exhibited predominantly constrained, non-directed movement in both D21 and T21 cells, consistent with recent studies demonstrating that centriolar satellites undergo largely diffusive rather than processive transport (Conkar et al., 2019). Although microtubules contribute to the spatial organization of centriolar satellites, the similar motility and exchange dynamics of PCM1 in D21 and T21 argue that altered transport does not account for its increased pericentrosomal accumulation in T21. We propose that elevated PCNT expands PCM1-rich pericentrosomal assemblies that act as capture sites for MIB1 and HERC2 during their transport to the centrosome. In this model, MIB1 and HERC2 continue to traffic to and accumulate at the centrosome, but their increased association with the expanded pericentrosomal PCM1 compartment reduces the efficiency of their centrosomal delivery. Because the centrosomal and pericentrosomal compartments draw upon a shared pool of proteins, even a modest elevation of PCNT can shift their steady-state distribution toward the pericentrosomal compartment without disrupting the underlying transport machinery. Thus, MIB1 and HERC2 are not excluded from the centrosome in T21, but their increased partitioning into pericentrosomal PCM1 assemblies reduces their centrosomal availability. Together, these findings suggest that centrosome function is governed not only by protein composition and transport but also by the spatial partitioning of proteins between centrosomal and pericentrosomal compartments.

This model raises the question of how the pericentrosomal compartment can accumulate more PCM1 and its associated proteins without altering their underlying dynamics. Fluorescence recovery after photobleaching (FRAP) revealed that despite the marked increase in pericentrosomal PCM1 in T21 cells, the mobile fraction and recovery kinetics of PCM1 were unchanged (Figure S4E), indicating that elevated PCNT does not increase the stability or lifetime of individual PCM1 interactions. Instead, both dynamically exchanging and stable or slowly exchanging PCM1 populations remained unchanged in T21. Although these data do not distinguish which population mediates pericentrosomal capture of MIB1 and HERC2, they support a model in which elevated PCNT remodels the pericentrosomal compartment by increasing its capacity to accommodate transient PCM1 interactions without fundamentally altering PCM1 exchange dynamics. Consequently, more PCM1, together with associated MIB1 and HERC2, can occupy the pericentrosomal compartment at steady state while remaining dynamically exchangeable. This increased compartment capacity provides a mechanism by which MIB1 and HERC2 can continue to reach the centrosome but do so less efficiently, delaying the accumulation of the machinery required for CP110 removal. Such dynamic partitioning is consistent with the delayed, rather than blocked, uncapping and ciliogenesis observed in T21 cells.

PCM1-positive centriolar satellites promote primary ciliogenesis by interacting with and regulating the localization of numerous centrosomal proteins (Hall et al., 2023; Odabasi et al., 2019; Prosser & Pelletier, 2020; Tollenaere et al., 2015; Wang et al., 2016). How centriolar satellites control centrosomal protein composition, however, remains poorly understood. Because centriolar satellite accumulation in T21 cannot be explained by altered transport, we asked whether the spatial distribution of PCM1 itself is the determinant. Overexpression or forced relocalization of PCM1 was sufficient to decrease MIB1 and HERC2 localization at the centrosome, demonstrating that the spatial distribution of PCM1 determines the availability of these ligases at the centrosome. Notably, increasing PCM1 abundance elevated PCM1 at both centrosomal and pericentrosomal regions yet reduced, rather than increased, centrosomal MIB1 or HERC2. The ubiquitin ligases instead accumulated within the expanded pericentrosomal PCM1 compartment, indicating that this compartment partitions MIB1 and HERC2 away from the centrosome even as the centrosomal PCM1 increases. Reducing PCM1 had the opposite effect and improved ciliogenesis in T21, consistent with excess pericentrosomal PCM1 acting as the limiting factor that restricts centrosomal availability of these ligases. That PCM1 reduction impairs ciliogenesis in disomic cells underscores that PCM1 is required for ciliogenesis but is detrimental in excess, and that its distribution rather than its abundance alone determines the outcome. PCM1 reduction also lowered centrosomal and pericentrosomal PCNT to disomic levels, indicating that PCM1 and PCNT reinforce one another’s accumulation. This reciprocal relationship may stabilize the expanded pericentrosomal architecture of T21 and further bias centriolar satellite-associated proteins away from the centrosome.

Timely removal of CP110 from the mother centriole is a critical licensing step in ciliogenesis and is coordinated by multiple pathways that regulate its ubiquitination and proteasomal degradation (Gonçalves et al., 2021; Nagai et al., 2018; Shen et al., 2021; Spektor et al., 2007; Tsang et al., 2008; Xie et al., 2023; Yadav et al., 2016). Although MIB1 and HERC2 are established mediators of CP110 ubiquitination and PCM1 has been implicated in their centrosomal localization, how the spatial organization of this pathway contributes to CP110 removal has remained unclear. Previous work showed that elevated PCNT in T21 produces pericentrosomal crowding that traps MYO5A and disrupts its accumulation at the mother centriole (Jewett et al., 2023), suggesting that increased PCNT can impair ciliogenesis by restricting the spatial access of ciliary factors to the centrosome. Our findings extend this model to the CP110 ubiquitination pathway, showing that elevated PCNT expands a PCM1-rich pericentrosomal compartment that redistributes MIB1 and HERC2 away from the mother centriole, thereby reducing the local availability of these E3 ligases to ubiquitinate CP110. Consistent with this model, T21 directly reduced CP110 ubiquitination, linking MIB1 and HERC2 mislocalization to the biochemical modification that initiates centriolar uncapping rather than to protein mislocalization alone. This disruption was selective because centrosomal TTBK2 was unchanged, while MIB1 and HERC2 were reduced, indicating that delayed uncapping results from selective disruption of CP110 ubiquitination rather than wholesale remodeling of the uncapping machinery. Importantly, impaired CP110 ubiquitination may act alongside other T21-associated defects in early ciliogenesis. Previous work showed that T21 reduced EHD1 accumulation at the mother centriole, which could further contribute to delayed uncapping by impairing the ciliary vesicle formation events that accompany CP110 removal (Jewett et al., 2023; Lu et al., 2026).Together, these findings identify spatial control of the CP110 ubiquitination machinery as a mechanism regulating centriolar uncapping and show that the timing of ciliogenesis is governed not only by the activity of CP110 regulators but also by their spatial access to the mother centriole.

Our findings provide a mechanistic framework linking altered centrosome organization to the ciliary defects observed in T21. T21 delays centriolar uncapping and the onset of ciliogenesis, revealing that the principal defect is kinetic rather than absolute. This distinction may be particularly important during development, when many cilia-dependent signaling pathways, including Hedgehog signaling, operate within tightly regulated temporal windows. Delayed cilium assembly could therefore alter the onset, duration, or magnitude of signaling without abolishing pathway activity, providing one explanation for how relatively modest defects in ciliogenesis contribute to the diverse developmental phenotypes of Down syndrome (Bergström et al., 2016; Haydar & Reeves, 2012; Richtsmeier et al., 2000). More broadly, remodeling of the pericentrosomal compartment by a 1.5-fold elevation of PCNT illustrates how a small change in gene dosage can alter centrosome organization and, through it, the timing of ciliogenesis and cilia-dependent signaling.

## SUPPLEMENTAL FIGURE LEGENDS

**Supplemental Figure 1 Trisomy 21 delays uncapping and ciliogenesis.** (**S1A**) Knockdown of PCNT reduces centrosomal and pericentrosomal PCNT in D21 and T21 24 hours post serum starvation. Quantification of centrosomal and pericentrosomal PCNT levels in D21 and T21 24 hours post serum starvation with or without knockdown of PCNT. Levels were normalized to D21 siC averages. Mean ± SD. *, p < 0.05. (**S1B**) Knockdown of CP110 reduces whole cell CP110 levels in D21 and T21 24 hours post serum starvation. Quantification of whole cell CP110 levels in D21 and T21 24 hours post serum starvation with or without knockdown of CP110. Levels were normalized to D21 siC or T21 siC controls. Mean ± SD. *, p < 0.05. (**S1C**) Knockdown of CP110 does not alter centrosomal and pericentrosomal PCNT levels in D21 and T21. Quantification of centrosomal and pericentrosomal PCNT levels in D21 and T21 24 hours post serum starvation with or without knockdown of CP110. Levels were normalized to D21 siC averages. Mean ± SD. *, p < 0.05. (**S1D**) Initiation of ciliogenesis with LPA inhibitor KI16245 phenocopies T21 ciliogenesis and centriolar uncapping delay. Left panel, quantification of centriolar uncapping frequency in D21 and T21 with or without LPA inhibition across an LPA inhibition time course. Mean ± SD. *, p < 0.05. Right panel, quantification of cilia frequency in D21 and T21 with or without LPA inhibition across an LPA inhibition time course. Mean ± SD. *, p < 0.05. (**S1E**) T21 increases centrosomal and pericentrosomal PCNT at every time point post serum starvation. Left panel, quantification of centrosomal PCNT in D21 and T21 across a serum starvation time course. Levels were normalized to D21 0 hour averages. Mean ± SD. *, p < 0.05. Right panel, quantification of pericentrosomal PCNT in D21 and T21 across a serum starvation time course. Levels were normalized to D21 0 hour averages. Mean ± SD. *, p < 0.05. (**S1F**) T21 increases centrosomal and pericentrosomal PCNT at every time point post LPA inhibition. Left panel, quantification of centrosomal PCNT in D21 and T21 across an LPA inhibition time course. Levels were normalized to D21 0 hour averages. Mean ± SD. *, p < 0.05. Right panel, quantification of pericentrosomal PCNT in D21 and T21 across an LPA inhibition time course. Levels were normalized to D21 0 hour averages. Mean ± SD. *, p < 0.05. (**S1G**) T21 increases total levels of CP110 and CEP97 across a serum starvation time course. Uncropped western blots depicting CP110 or CEP97 levels (magenta) with respective loading controls (green) across a serum starvation time course.

**Supplemental Figure 2 Trisomy 21 reduces CP110 ubiquitination and centrosomal localization of E3 ligases.** (**S2A**) T21 increases whole cell levels of MIB1. Left panel, quantification of whole cell MIB1 in D21 and T21 0.5 hours post serum starvation. Levels were normalized to D21 averages. Mean ± SD. *, p < 0.05. Right panel, western blot of MIB1 levels (red) in D21 and T21 0.5 hours post serum starvation with Lamin A/C loading control (green). (**S2B**) T21 increases whole cell levels of HERC2. Quantification of whole cell HERC2 in D21 and T21 0.5 hours post serum starvation. Levels were normalized to D21 averages. Mean ± SD. *, p < 0.05. (**S2C**) T21 does not alter centrosomal TTBK2 levels. Quantification of centrosomal TTBK2 in D21 and T21 0.5 hours post serum starvation. Levels were normalized to D21 averages. Mean ± SD. *, p < 0.05. (**S2D**) T21 reduces CP110 ubiquitination. Uncropped western blots of CP110 (magenta) and respective loading controls (green) in D21 and T21 after immunoprecipitation (IP) of ubiquitin with or without the presence of proteasomal inhibitor MG132. (**S2E**) Uncropped western blots of ubiquitin and CP110 in D21 and T21 after immunoprecipitation (IP) of CP110 with or without the presence of proteasomal inhibitor MG132.

**Supplemental Figure 3 Trisomy 21 increases PCNT and PCM1 pericentrosomal accumulation during centriolar uncapping.** (**S3A**) MIB1 and HERC2 colocalize with PCM1 in D21 and T21 0.5 hours post serum starvation. Left panels, cells stained for MIB1 (grayscale) and PCM1 (green) or HERC2 (grayscale) and PCM1 (green). Yellow arrows denote colocalized puncta. Scale bar, 5 µm. Right panels, quantification of Pearson correlation coefficients for PCM1 and MIB1 or PCM1 and HERC2 0.5 hours post serum starvation in D21 and T21. Mean ± SD. *, p < 0.05. (**S3B**) T21 increases centrosomal and pericentrosomal PCM1 across a serum starvation or LPA inhibition time course. Levels were normalized to D21 0 hour averages. Mean ± SD. *, p < 0.05. (**S3C**) Knockdown of CP110 reduces centrosomal and pericentrosomal PCM1 levels in D21 and T21 0.5 hours post serum starvation. Quantification of centrosomal and pericentrosomal PCM1 in D21 and T21 with or without knockdown of CP110. Levels were normalized to D21 siC averages. Mean ± SD. *, p < 0.05. (**S3D**) Knockdown of PCM1 rescues cilia frequency in T21. Left panel, quantification of centrosomal and pericentrosomal PCM1 with or without knockdown of PCM1 in D21 and T21 24 hours post serum starvation. Levels were normalized to D21 siC averages. Mean ± SD. *, p < 0.05. Middle panel, quantification of cilia frequency in D21 and T21 with or without knockdown of PCM1 24 hours post serum starvation. Mean ± SD. *, p < 0.05. Right panel, quantification of centrosomal and pericentrosomal PCNT with or without knockdown of PCM1 24 hours post serum starvation. Levels were normalized to D21 siC averages. Mean ± SD. *, p < 0.05.

**Supplemental Figure 4 Trisomy 21 does not alter PCNT or PCM1 motility.** (**S4A**) Overexpressing PCM1 with low levels of doxycycline maintains D21 centrosomal PCNT levels at 1X. Left panel, quantification of whole cell PCM1 levels in D21 across doxycycline titration. Levels were normalized to the D21 1 µg/mL average. Mean ± SD. Right panel, quantification of D21 centrosomal PCNT levels across doxycycline titration. Levels were normalized to D21 no doxycycline average. Mean ± SD. (**S4B**) PCNT-mNeon localizes to centrosomal and pericentrosomal regions. Cell expressing PCNT-mNeon. Scale bar, 5 µm. (**S4C**) T21 does not alter average PCNT puncta motility. Top panels, plots showing average PCNT track trajectories in the pericentrosomal region 0.5 hours post serum starvation. Trajectories are colored by average speed (µm/second). Middle panels, T21 does not alter PCNT puncta trajectory direction. Plots showing average PCNT track trajectories in the pericentrosomal region 0.5 hours post serum starvation. Trajectories are colored by direction of movement either toward or away from the centrosome. Bottom panels, T21 does not alter PCNT MSD. Individual MSD trajectories for PCNT puncta in D21 and T21 0.5 hours post serum starvation. (**S4D**) Top panel, average PCNT foci velocities in D21 and T21. Middle panel, PCNT foci directionality ratios in D21 and T21. Bottom panel, PCNT MSD slopes for D21 and T21. Mean ± SD. *, p < 0.05. (**S4E**) T21 does not alter PCNT or PCM1 pericentrosomal recovery 0.5 hours post serum starvation. Left panels, representative images depicting PCNT-mNeon or PCM1-GFP recovery across a photobleaching time course. Scale bar, 5 µm. Right panels, quantification of PCNT-mNeon or PCM1-GFP fluorescence recovery after photobleaching in D21 and T21 0.5 hours post serum starvation. Mean ± SD.

**Supplemental Figure 5 Elevated PCM1 mislocalizes MIB1 and HERC2 and causes uncapping and ciliogenesis defects**. (**S5A**) PCM1 overexpression reduces centriolar uncapping and ciliogenesis in D21 with or without knockdown of PCNT. Left panel, relative change in ciliation 24 hours post serum starvation in D21 cells with or without knockdown of PCNT. Mean ± SD. *, p < 0.05. Right panel, relative change in uncapping 24 hours post serum starvation in D21 cells with or without knockdown of PCNT. Mean ± SD. *, p < 0.05. (**S5B**) PCM1 overexpression reduces centriolar uncapping and ciliogenesis in T21. Left panels, quantification of cilia frequency and relative change in ciliation 24 hours post serum starvation in T21 cells with or without knockdown of PCNT. Mean ± SD. *, p < 0.05. Right panel, quantification of uncapping frequency and relative change in uncapping 24 hours post serum starvation in T21 cells with or without knockdown of PCNT. Mean ± SD. *, p < 0.05. (**S5C**) PCM1 overexpression reduces centrosomal MIB1 and HERC2 levels in T21 0.5 hours post serum starvation. Left panels, quantification of centrosomal MIB1 levels and relative change in MIB1 0.5 hours post serum starvation in T21 cells with or without knockdown of PCNT. Levels were normalized to D21 siC averages. Mean ± SD. *, p < 0.05. Right panels, quantification of centrosomal HERC2 levels and relative change in HERC2 0.5 hours post serum starvation in T21 cells with or without knockdown of PCNT. Levels were normalized to D21 siC averages. Mean ± SD. *, p < 0.05. (**S5D**) PCM1 overexpression increases pericentrosomal HERC2 levels in T21 0.5 hours post serum starvation with knockdown of PCNT. Quantification of pericentrosomal MIB1 and HERC2 levels 0.5 hours post serum starvation in T21 cells with knockdown of PCNT. Levels were normalized to D21 siC averages. Mean ± SD. *, p < 0.05. (**S5E**) Knockdown of PCNT does not alter the formation of PCM1 assemblies in PCM1 overexpression contexts. D21 cells expressing PCM1-GFP stained for CEP192 (grayscale) with or without knockdown of PCNT.(**S5F**) Altering PCM1 localization pharmacologically mislocalizes MIB1 and HERC2. Top panel images, D21 cells stained for MIB1 (grayscale) and PCM1 (green) or HERC2 (grayscale) and PCM1 (green) 24 hours post serum starvation and MG132 treatment. Scale bar, 5 µm. Top panel graphs, quantification of centrosomal PCM1, MIB1, or HERC2 levels in D21 and T21 with or without MG132 treatment. Levels were normalized to D21-MG132 averages. Mean ± SD. *, p < 0.05. Bottom panel images, D21 cells stained for MIB1 (grayscale) and PCM1 (green) or HERC2 (grayscale) and PCM1 (green) 24 hours post serum starvation and nocodazole (NZ) treatment. Scale bar, 5 µm. Bottom panel graphs, quantification of centrosomal PCM1, MIB1, or HERC2 levels in D21 and T21 with or without nocodazole treatment. Levels were normalized to D21-NZ averages. Mean ± SD. *, p < 0.05. (**S5G**) Rapamycin does not alter background PCNT levels or cilia frequency. Left panel, centrosomal and pericentrosomal PCNT levels in D21 and T21 cells with or without rapamycin. Levels were normalized to D21 - RAPA averages. Right panel, D21 and T21 cilia frequencies with or without rapamycin. Mean ± SD. *, p < 0.05. (**S5H**) Pulling PCM1 to or away from the centrosome does not alter centrosomal PCNT levels. Left panel, D21 and T21 centrosomal PCNT levels in cells overexpressing PCM1-FKBP and BICD2-FRB with or without rapamycin. Levels were normalized to D21-RAPA averages. Right panel, D21 and T21 centrosomal PCNT levels in cells overexpressing PCM1-FKBP and LYN11-FRB with or without rapamycin. Levels were normalized to D21-RAPA averages. Mean ± SD. *, p < 0.05.

## ACKNOWLEDGEMENTS

We would like to thank Joaquin Espinosa, Molishree Joshi for her work in generating the endogenously tagged PCNT-mNeon D21 and T21 cell lines, Kelly Sullivan, Elif Firat-Karalar for the PCM1-GFP-FKBP and BICD2-HA-FKB constructs, Andrew Neumann and Rytis Prekeris for assistance with the Signal-Seeker Ubiquitination Detection Kit, and the Pearson lab for helpful discussions. This research was funded by NIH/NIGMS R35GM140813 to C.G.P., the Bolie Fellowship to B.L.M., the Linda Crnic Institute for Down Syndrome, and the Global Down Syndrome Foundation.

## METHODS

### Key resources table

| REAGENT or RESOURCE | SOURCE | IDENTIFIER |
| --- | --- | --- |
| **Antibodies** |  |  |
| Mouse monoclonal anti-Polyglutamylated Tubulin (GT335) | Adipogen | RRID: AB_2490210 |
| Rabbit polyclonal anti-CP110 | Proteintech | RRID: AB_10638480 |
| Mouse monoclonal anti-Alpha tubulin (DM1A) | Sigma | RRID: AB_2490210 |
| Mouse monoclonal anti-LAMIN A/C | Cell Signaling | RRID: AB_10545756 |
| Mouse monoclonal anti-HERC2 | BD Biosciences | RRID: AB_399728 |
| Rabbit polyclonal anti-MIB1 | Millipore Sigma | RRID: AB_1841007 |
| Rabbit polyclonal anti-CEP97 | Proteintech | RRID: AB_11182378 |
| Rabbit polyclonal anti-PCNT | Abcam | RRID: AB_304461 |
| Mouse monoclonal anti-ubiquitin | Santa Cruz | RRID: AB_62842 |
| Rabbit polyclonal anti-PCM1-488 | Proteintech | RRID: AB_2919167 |
| Rabbit polyclonal anti-ARL13B | Proteintech | RRID: AB_2060867 |
| Goat polyclonal anti-CEP192 | Andrew Holland Lab | NA |
| Mouse monoclonal anti-ARL13B | DSHB | RRID: AB_11000053 |
| Rabbit polyclonal anti-PCM1 | Cell Signaling | RRID: AB_10556960 |
| Mouse monoclonal anti-HA (16B12) | Biolegend | RRID: AB_2565336 |
| Rabbit polyclonal anti-TTBK2 | Proteintech | RRID: AB_2211507 |
| Mouse monoclonal anti-rabbit IgG | Cell Signaling | RRID: AB_1549606 |
| Donkey anti-rabbit Alexa 594 | Invitrogen | RRID: AB_141637 |
| Donkey anti-goat Alexa 594 | Invitrogen | RRID: AB_142540 |
| Donkey anti-rabbit Alexa 488 | Invitrogen | RRID: AB_2535792 |
| Donkey anti-goat Alexa 633 | Invitrogen | RRID: AB_14149 |
| Donkey anti-mouse Alexa 647 | Invitrogen | RRID: AB_162542 |
| IRDye 680RD goat anti-rabbit | LiCor | RRID: AB_10956166 |
| IRDye 800CW goat anti-mouse | LiCor | RRID: AB_621842 |
| **Biological samples** |  |  |
| hTERT immortalized RPE1 D21 | Lane et al., 2014 | NA |
| hTERT immortalized RPE1 T21 | Lane et al., 2014 | NA |
| **Chemicals, peptides, and recombinant proteins** |  |  |
| Bovine serum albumin | Sigma | A3912 |
| Citifluor AF1 | Electron Microscopy Sciences | 17970-100 |
| Collagen | Sigma | C9791 |
| DMEM:F12 | Gibco | 11320-033 |
| Doxycycline | Sigma Aldrich | 17086-28-1 |
| FBS | Pheonix Scientific | PS-100 |
| Fluorobrite DMEM | Invitrogen | A1896701 |
| Hoescht 33258 | Invitrogen | H3569 |
| Igepal | Sigma | 18896 |
| KI16245 | Millipore Sigma | SML0971-5MG |
| Lipofectamine 2000 | Invitrogen | 11668027 |
| Lipofectamine RNAi Max | Invitrogen | 13778030 |
| Methanol | Thermostat | L13255.0F |
| MG132 | Millipore Sigma | C2211-5MG |
| Nocadazole | Sigma Aldrich | SML1665-1ML |
| Opti-MEM | Gibco | 31985-070 |
| Rapamycin | Med Chem Express | 53123-88-9 |
| Triton-X-100 | Sigma Aldrich | T9284-500ML |
| Trypsin | Gibco | 15090-046 |
| Tween 20 | Millipore Sigma | P9416-50ML |
| **Critical commercial assays** |  |  |
| Immunoprecipitation Kit - Dynabeads Protein G | Invitrogen | 10007D |
| Signal-Seeker Ubiquitination Detection Kit | Cytoskeleton | BK161-S |
| **Experimental Models: Cell Lines** |  |  |
| RPE1 D21 PCNT mNeon | This study | NA |
| RPE1 T21 PCNT mNeon | This study | NA |
| RPE1 D21 PCM1-FKBP-GFP | This study | NA |
| RPE1 T21 PCM1-FKBP-GFP | This study | NA |
| RPE1 D21 PCM1-FKBP-GFP BICD2-FRB-HA | This study | NA |
| RPE1 T21 PCM1-FKBP-GFP BICD2-FRB-HA | This study | NA |
| RPE1 D21 PCM1-FKBP-GFP LYN11-FRB-RFP | This study | NA |
| RPE1 T21 PCM1-FKBP-GFP LYN11-FRB-RFP | This study | NA |
| **Oligonucleotides** |  |  |
| Human Pericentrin siRNA (Smart Pool) | Dharmacon | M-012172-01-0005 |
| Mission siRNA Universal Negative Control | Sigma | SIC001-10NMOL |
| siPCM1 A (On-target plus siRNA) | Dharmacon | J-005165-07-0002 |
| siPCM1 B (On-target plus siRNA) | Dharmacon | J-005165-09-0002 |
| CP110 custom siRNA (on-target plus siRNA) | Dharmacon | SO-3052251G |
| **Recombinant DNA** |  |  |
| pCW-57.1-PCM1-FKBP-GFP | This study | NA |
| pCW-57.1-LYN11-FRB-RFP | This study | NA |
| pCW-57.1-BICD2-FRB-HA | This study | NA |
| **Software and algorithms** |  |  |
| Slidebook | 3i | https://www.intelligent-imaging.com/slidebook |
| FIJI | LOCI/Wisconsin-Madison | RRID: SCR_002285 |
| ImageJ | Wayne Rhasband/NIH | RRID: SCR_003070 |
| TrackMate (ImageJ plugin) | ImageJ | https://imagej.net/plugins/trackmate/ |
| MatLab R2023b | MathWorks | https://www.mathworks.com/products/new_products/release2023b.html |
| Excel 365 | Microsoft | https://www.office.com/ |
| Graphpad Prism | Graphpad | https://www.graphpad.com/features |
| Coloc2 (ImageJ plugin) | ImageJ | https://imagej.net/plugins/coloc-2 |
| **Other** |  |  |
| Nikon Eclipse Ti-E | Nikon |  |
| 100X Plan Apochromat Objective (NA 1.40) | Nikon | 01901 |
| Teledyne Photometrics Kinetix 22 cMOS Camera | Teledyne | 01-KINETIX-22MM-M-C |
| Nikon Eclipse Ti | Nikon |  |
| Teledyne Photometrics Prime95B sCMOS Camera | Teledyne |  |
| CSU-X1 Spinning Disk | Nikon |  |
| OkoLab Controlled Environment Stage | OkoLab |  |
| OkoLab H301-K-Frame | OkoLab |  |
| 60X Plan Apochromat Objective (NA 1.40) | Nikon |  |
| 1.5 glass coverslips | Electron Microscopy Sciences | 72230-01 |

### Cell Culture

Isogenic disomy 21 (D21) and trisomy 21 (T21) hTERT-immortalized retinal pigment epithelial (RPE-1) cell lines were generated as previously described (Lane et al., 2014). Cells were maintained in DMEM/F12 supplemented with 10% fetal bovine serum and 1% penicillin-streptomycin in a humidified incubator at 37°C with 5% CO₂. Cells were passaged at approximately 80–90% confluency using 0.25% trypsin at a 1:10 split ratio. Primary ciliogenesis was induced by serum depletion in DMEM/F12 containing 0.5% fetal bovine serum and 1% penicillin-streptomycin for the indicated experimental times, or by treatment with 10 μM of the LPA receptor inhibitor KI16245 for the indicated experimental times. All cell culture reagents are listed in the Key Resources Table. Cell lines were routinely tested for mycoplasma contamination every six months and were confirmed to be mycoplasma-free prior to experiments.

### Immunofluorescence

Twelve-millimeter glass coverslips were cleaned by incubation in 1 M HCl at 50°C for 16 hours, followed by sequential sonication for 30 minutes each in water, 50%, 70%, and 95% ethanol. Coverslips were coated with collagen, air dried for 15 minutes, and UV crosslinked for an additional 15 minutes before washing three times with phosphate-buffered saline (PBS; 1 mM KH₂PO₄, 155 mM NaCl, 3 mM Na₂HPO₄·7H₂O, pH 7.4). Cells were seeded onto prepared coverslips and allowed to adhere overnight. Cells were fixed in ice-cold 100% methanol for 10 minutes at −20°C, permeabilized in 0.5% Triton X-100 in PBS for 10 minutes at room temperature, and washed three times in PBS. Cells were blocked for 1 hour at room temperature in PBS containing 0.5% bovine serum albumin (BSA), 0.5% IGEPAL (NP-40), 1 mM MgCl₂, and 1 mM NaN₃. Primary antibodies were diluted in blocking buffer and incubated for 1 hour at room temperature, followed by three 5-minute washes in PBS. Alexa Fluor-conjugated secondary antibodies and Hoechst 33258 (1 μg/mL) were diluted in blocking buffer and incubated for 1 hour at room temperature. Coverslips were washed four times for 5 minutes in PBS, mounted using Citifluor AF1 mounting medium, and sealed with clear nail polish. Antibodies are listed in the Key Resources Table.

### Live-cell Imaging

Live-cell imaging was performed using custom glass-bottom imaging dishes prepared by replacing the bottom of 35-mm tissue culture dishes with 30-mm glass coverslips affixed using aquarium-grade silicone sealant. Following curing and UV sterilization, dishes were coated with collagen overnight at 4°C, washed with PBS, and seeded with cells. Prior to imaging, culture medium was replaced with FluoroBrite DMEM supplemented with 1% penicillin-streptomycin. Cells expressing endogenous PCNT-mNeon or exogenous PCM1-FKBP-GFP were serum starved for 30 minutes before imaging. Time-lapse images were acquired on a Nikon Eclipse Ti-E microscope equipped with a 100× Plan Apochromat objective (NA 1.40), a Teledyne Photometrics Kinetix22 sCMOS camera, and an Okolab environmental chamber maintained at 37°C with 5% CO₂. Images were acquired at 1-second intervals for 120 seconds using identical acquisition settings for all experiments.

### Particle Tracking

Time-lapse image sequences were imported into FIJI/ImageJ and analyzed using the TrackMate plugin. PCNT-mNeon and PCM1-FKBP-GFP puncta were automatically detected using the Laplacian of Gaussian (LoG) detector and linked into trajectories using the Linear Assignment Problem (LAP) tracker. Detection and tracking parameters were maintained for all datasets within an experiment. Tracks corresponding to puncta present for fewer than 8 frames or exhibiting obvious tracking artifacts were excluded from subsequent analyses. TrackMate XML files containing puncta coordinates and trajectory information were exported for downstream analysis in MATLAB.

### MATLAB Analysis

Custom MATLAB scripts used for trajectory analysis, mean squared displacement analysis, directionality analysis, and puncta density quantification are available at https://github.com/baileylmccurdy2026/foci-puncta-analysis/tree/main. Portions of the code were developed with assistance from OpenAI ChatGPT and subsequently validated by the authors. TrackMate XML files were imported into MATLAB (MathWorks R2023b) using custom analysis scripts. Particle trajectories were reconstructed from the exported x-, y-, z-coordinate and frame information and converted to physical units using the TrackMate frame interval and pixel calibration. Instantaneous velocities were calculated from frame-to-frame displacement of individual puncta. Mean squared displacement (MSD) was calculated for each trajectory from two-dimensional coordinates across increasing lag times, and the diffusion coefficient was estimated from the slope of the linear fit. Motion type was assessed by fitting the MSD on a log-log scale to determine the anomalous diffusion exponent (α). Trajectories were classified as constrained (α < 1), diffusive (α ≈ 1), or directed (α > 1). Trajectory directionality was quantified using custom MATLAB scripts by calculating the change in distance of each punctum relative to the centroid of all tracked puncta. Linear regression of punctum distance from the centroid over time was used to classify trajectories as moving toward (negative slope) or away from (positive slope) the centrosome. Pericentrosomal puncta coordinates exported from FIJI were analyzed using custom MATLAB scripts. Coordinates from all cells within an experimental condition were combined to calculate a global centroid. Because all trajectories were centered on the centrosome prior to export from FIJI, the global trajectory centroid corresponded to the centrosome and was used as the reference point for directionality calculations. Local puncta density was calculated using a fixed-radius nearest-neighbor search, and marker size was scaled according to local puncta density. Puncta color was determined by normalized distance from the global centroid, allowing simultaneous visualization of puncta density and radial distribution within the pericentrosomal compartment. Radial distribution analyses were additionally performed by assigning puncta to normalized distance bins relative to the centroid and calculating puncta counts within each radial interval.

### Anchor-Away

To manipulate PCM1 localization, RPE-1 D21 and T21 cells stably expressing PCM1-FKBP-GFP together with either BICD2-FRB-HA or LYN11-FRB-RFP were used. Expression of pCW57.1-derived constructs was induced by treatment with 1 μg/mL doxycycline. Primary ciliogenesis was induced by serum starvation in DMEM/F12 supplemented with 0.5% fetal bovine serum and 1% penicillin-streptomycin. Simultaneously with serum starvation, cells were treated with 1ug/mL doxycycline and 500 nM rapamycin to induce heterodimerization of FKBP- and FRB-tagged proteins. Rapamycin-mediated recruitment of PCM1-FKBP-GFP to BICD2-FRB-HA relocalized PCM1 to the centrosome, whereas recruitment to LYN11-FRB-RFP redirected PCM1 to the plasma membrane. Cells were fixed 24 hours after serum starvation, doxycycline induction, and rapamycin treatment for immunofluorescence analysis.

### RNAi

siRNA transfections were performed using Lipofectamine RNAiMAX according to the manufacturer’s instructions. siRNAs were used at a final concentration of 25 nM in DMEM/F12 supplemented with 10% fetal bovine serum. MISSION siRNA Universal Negative Control #1 was used as the non-targeting control. Cells were incubated with siRNA complexes for 8 hours, after which transfection medium was replaced with fresh growth medium overnight. The following day, primary ciliogenesis was induced by serum depletion in DMEM/F12 containing 0.5% fetal bovine serum for the indicated times prior to fixation. PCNT, PCM1, and CP110 knockdowns were confirmed by immunofluorescence or immunoblotting prior to quantitative analysis. All siRNAs are listed in the Key Resources Table.

### Generation of cell lines

Stable PCM1-FKBP-GFP cell lines were generated first and subsequently transduced with either BICD2-FRB-HA or LYN11-FRB-RFP to establish dual-expression anchor-away cell lines. Endogenously tagged PCNT-mNeon D21 and T21 RPE-1 cell lines used for live-cell imaging were generated previously by CRISPR/Cas9-mediated endogenous tagging of the PCNT locus (McCurdy et al., 2022). Lentiviral particles were produced in HEK293T cells by co-transfecting expression constructs with the packaging plasmids psPAX2 and pMD2.G using Lipofectamine 2000. Viral supernatants were collected 24 and 48 hours after transfection and used to transduce target cells in the presence of 2 μg/mL polybrene. For cell lines expressing multiple constructs, sequential rounds of lentiviral transduction and antibiotic selection were performed to establish stable polyclonal cell populations. Transduced cells were selected with either 10 μg/mL puromycin for 10 days or 10 μg/mL blasticidin for 14 days following each round of transduction. Stable cell lines harboring pCW57.1-derived constructs were induced with doxycycline overnight prior to experimentation. Fixed-cell experiments were performed following induction with 1 μg/mL doxycycline. For live-cell imaging of PCM1-FKBP-GFP, cells were induced with 8 ng/mL doxycycline, a concentration selected to provide sufficient fluorescence while avoiding detectable changes in endogenous PCNT abundance.

### Fluorescence recovery after photobleaching (FRAP)

Live-cell FRAP experiments were performed using Nikon Eclipse Ti-E microscope equipped with a controlled environment stage (OkoLab; H301-K-Frame; 37°C; 5% CO_2_) and an Okolab environmental chamber maintained at 37°C with 5% CO₂. PCNT-mNeon and PCM1-FKBP-GFP puncta were photobleached using a 488-nm laser. Three pre-bleach images were acquired at 1-second intervals, followed by photobleaching and a single immediate post-bleach image. Fluorescence recovery was monitored every 5 seconds for 5 minutes. Fluorescence intensity was quantified and normalized to minimum intensity (at time of photobleaching) and the maximum intensity (prior to photobleaching) within each acquisition. Recovery curves were averaged across cells and plotted as normalized fluorescence intensity over time.

### Fluorescence microscopy

Widefield fluorescence imaging and fluorescence recovery after photobleaching (FRAP) were performed using a Nikon Eclipse Ti-E microscope equipped with a 100× Plan Apochromat objective (NA 1.40) and a Teledyne Photometrics Kinetix22 sCMOS camera. Live-cell imaging was performed using an Okolab environmental chamber maintained at 37°C with 5% CO₂. Confocal fluorescence imaging was performed using a Nikon Eclipse Ti inverted microscope equipped with a CSU-X1 (Yokogawa) spinning disk confocal unit, a 60× Plan Apochromat objective (NA 1.40) or a 100× Plan Apochromat objective (NA 1.40), and a Teledyne Photometrics Prime95B sCMOS camera. All image acquisition was performed using SlideBook 6 software (Intelligent Imaging Innovations). Identical laser power, exposure times, camera settings, and image processing parameters were maintained for all images compared quantitatively within an experiment.

### Fluorescence quantification

Primary cilia were identified by ARL13B-positive axonemes extending from CEP192-labeled centrosomes. Cells were scored as uncapped when the mother centriole lacked detectable CP110 staining and was identified using CEP192 or GT335 labeling. For all fluorescence intensity measurements, centrosomes were identified by the centroid of CEP192 staining. Centrosomal fluorescence was quantified using a circular region of interest (ROI) with a 1.2-μm radius centered on the CEP192 centroid. Pericentrosomal fluorescence was quantified using an annular ROI extending from 1.2 to 5 μm from the centrosome. Mean fluorescence intensity was measured from maximum-intensity projected z-stacks in FIJI/ImageJ. Local background was determined from the average intensity of three intracellular regions lacking specific staining and subtracted from each measurement. Fluorescence intensities were normalized to the average value of the untreated D21 control for each independent experiment before pooling biological replicates. Pericentrosomal puncta were quantified from maximum-intensity projected z-stacks using the *Find Maxima* function in FIJI/ImageJ. Local intensity maxima were identified using a constant noise tolerance threshold for all images within an experiment. Detected puncta were assigned to concentric annular regions centered on the CEP192 centroid, and puncta number was quantified within each radial bin within the entire pericentrosomal ROI (1.2–5 μm), as indicated. Colocalization between PCM1 and PCNT, MIB1, or HERC2 was quantified using the Coloc2 plugin in FIJI/ImageJ. Pearson’s correlation coefficients were calculated from maximum-intensity projected z-stacks within centrosomal or pericentrosomal regions of interest. Identical acquisition settings and analysis parameters were maintained for all images within an experiment. To visualize average fluorescence distributions, maximum-intensity projected images from individual cells were spatially aligned and combined into an average intensity projection using FIJI/ImageJ. Average images were pseudocolored using the Fire lookup table (LUT) to generate representative fluorescence heat maps.

### Immunoprecipitation

To assess CP110 ubiquitination, cells were serum starved in DMEM/F12 supplemented with 0.5% fetal bovine serum and 1% penicillin-streptomycin in the presence of 10 μM MG132 for 4 hours prior to lysis. For ubiquitin pull-downs, ubiquitinated proteins were enriched using the Signal-Seeker Ubiquitination Detection Kit (Cytoskeleton) according to the manufacturer’s instructions. The enriched ubiquitinated protein fraction was then analyzed by SDS-PAGE and immunoblotting for CP110 to assess ubiquitinated CP110. For reciprocal immunoprecipitation, cell lysates were incubated with anti-CP110 antibody and Dynabeads Protein G (Invitrogen) according to the manufacturer’s instructions to immunoprecipitate CP110. Immunoprecipitated CP110 was subsequently analyzed by immunoblotting with anti-ubiquitin antibodies to detect ubiquitinated CP110.

### Immunoblotting

Cells were washed three times with ice-cold PBS, harvested using a rubber cell scraper, and pelleted by centrifugation. Cell pellets were lysed for 1 hour on ice in lysis buffer containing 50 mM Tris-HCl (pH 7.4), 150 mM NaCl, 1% NP-40, 0.5% sodium deoxycholate, and freshly added protease inhibitor cocktail. Lysates were clarified by centrifugation at maximum speed for 30 min at 4°C, and protein concentrations were equalized prior to denaturation in SDS sample buffer. Equal amounts of protein or immunoprecipitated samples were resolved by SDS-PAGE on 10% polyacrylamide gels and transferred to PVDF membranes. Membranes were blocked overnight at 4°C in Tris-buffered saline (TBS; 20 mM Tris, 150 mM NaCl, pH 7.6)containing 0.05% Tween-20 and 5% nonfat dry milk before incubation with primary antibodies overnight at 4°C. Following washing in PBS containing 0.05% Tween-20, membranes were incubated with IRDye 680RD goat anti-rabbit or IRDye 800CW goat anti-mouse secondary antibodies (LI-COR) for 1 hour at room temperature. Immunoblots were imaged using a LI-COR Odyssey imaging system, and band intensities were quantified in FIJI/ImageJ.

### Pharmacological manipulation of PCM1 localization

To alter PCM1 localization pharmacologically, cells were serum starved in DMEM/F12 supplemented with 0.5% fetal bovine serum and treated with either MG132 or nocodazole. Aggresome formation was induced by treatment with 10 μM MG132 for 24 hours during serum starvation. Centriolar satellite dispersal was induced by treatment with 0.1 μM nocodazole for 24 hours during serum starvation to depolymerize microtubules. Following treatment, cells were fixed and processed for immunofluorescence as described above.

### Statistical Tests

Data were analyzed using Microsoft Excel and GraphPad Prism. All experiments were performed with a minimum of three independent biological replicates unless otherwise indicated (Table 1). For fluorescence microscopy experiments, each biological replicate consisted of an independent cell culture. Quantitative data are presented as mean ± standard deviation (SD). Statistical significance between two groups was assessed using unpaired, two-tailed Student’s *t* tests. Exact sample sizes (n) are reported in Table 1. Differences were considered statistically significant if *p* < 0.05 (Table 1).

## ABREVIATIONS

PCNT: Pericentrin
PCM: Pericentriolar material
MT: Microtubule
MTOC: Microtubule organizing center
D21: Disomy 21
T21: Trisomy 21
MSD: Mean squared displacement
FRAP: Fluorescence recovery after photobleaching

## Notes

### Competing Interest Statement

The authors have declared no competing interest.

