## Supplementary figures and images for "Aberrant PCM1 accumulation in Trisomy 21 mislocalizes E3 ligases, delaying primary ciliogenesis"

### Supplemental Figures

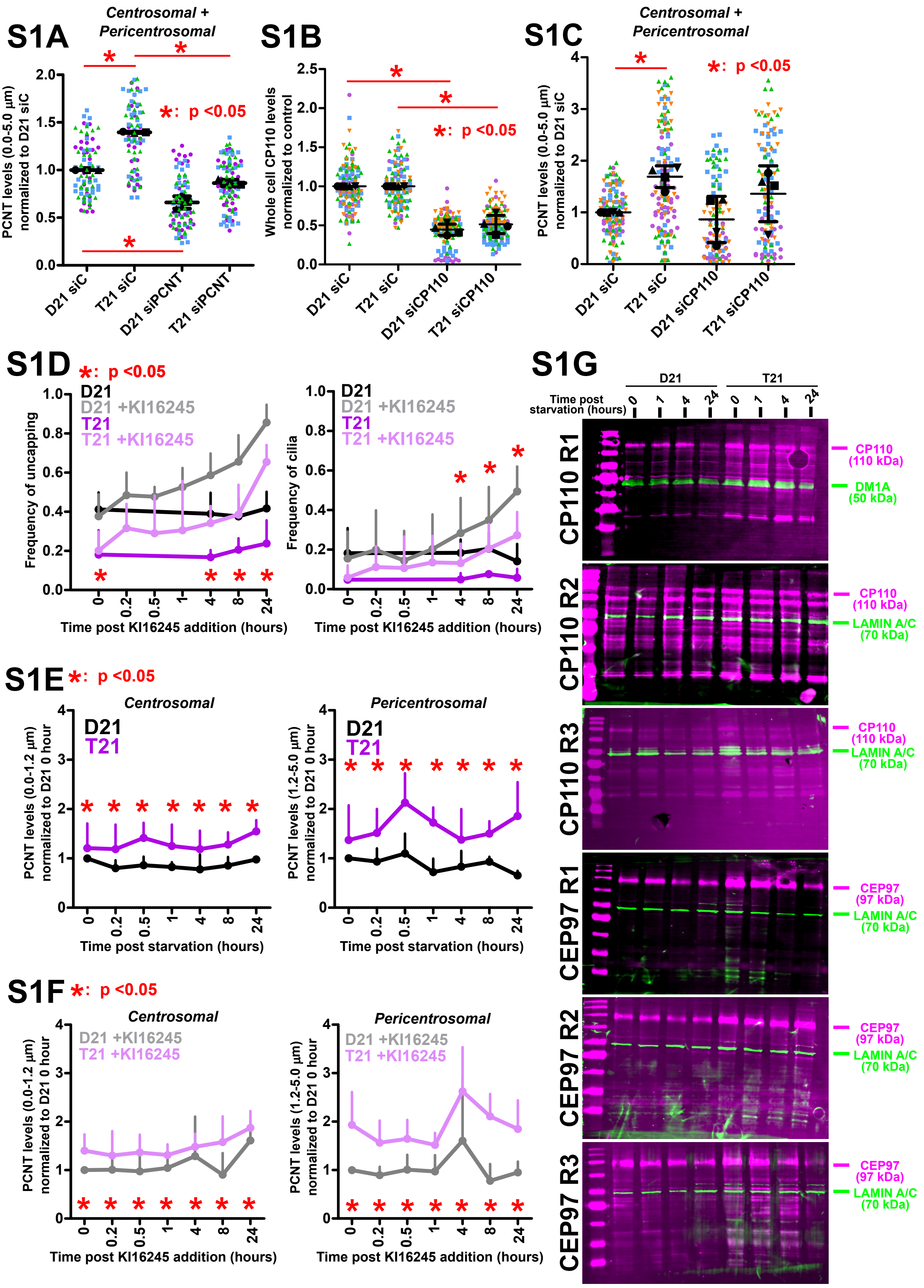

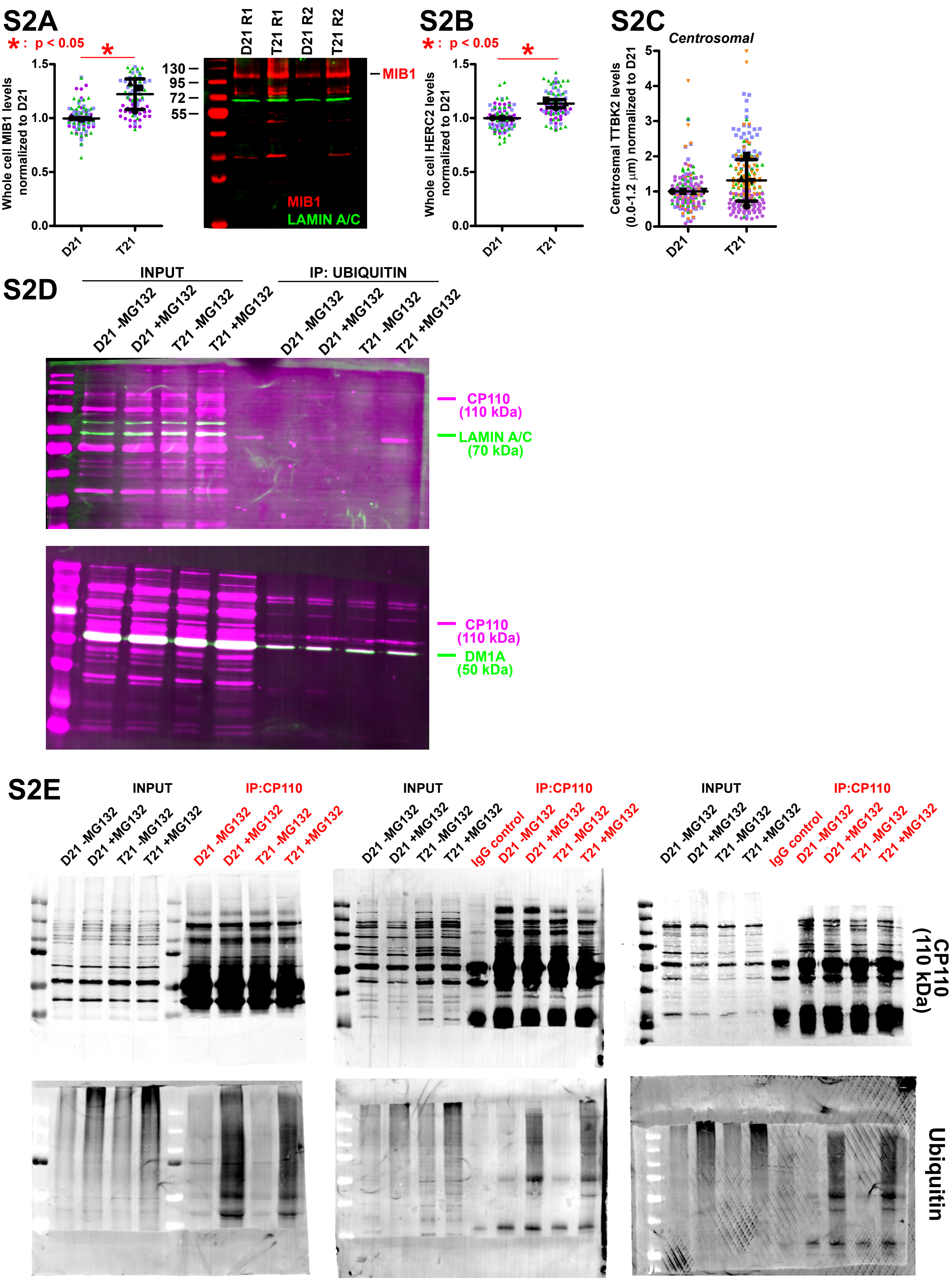

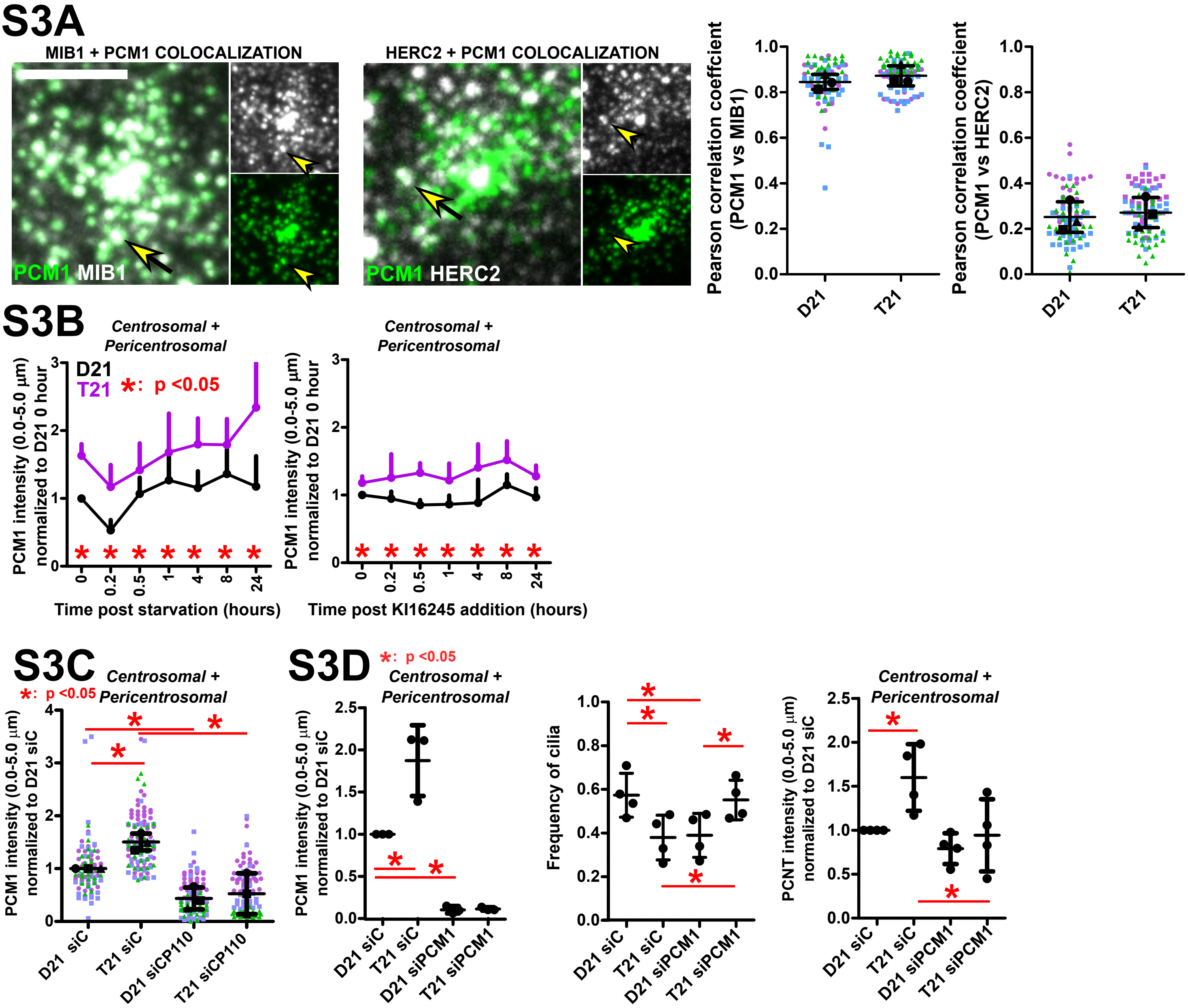

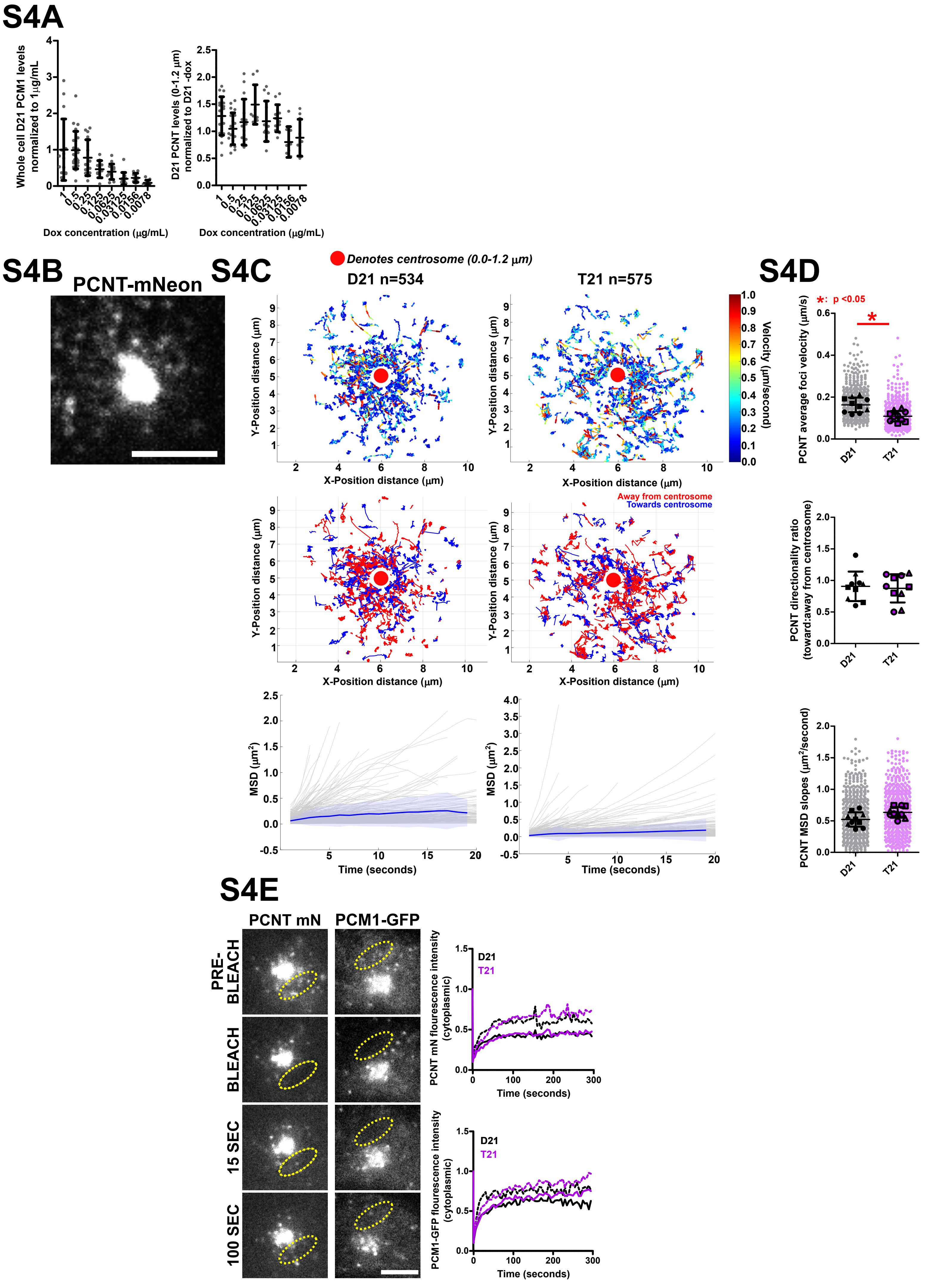

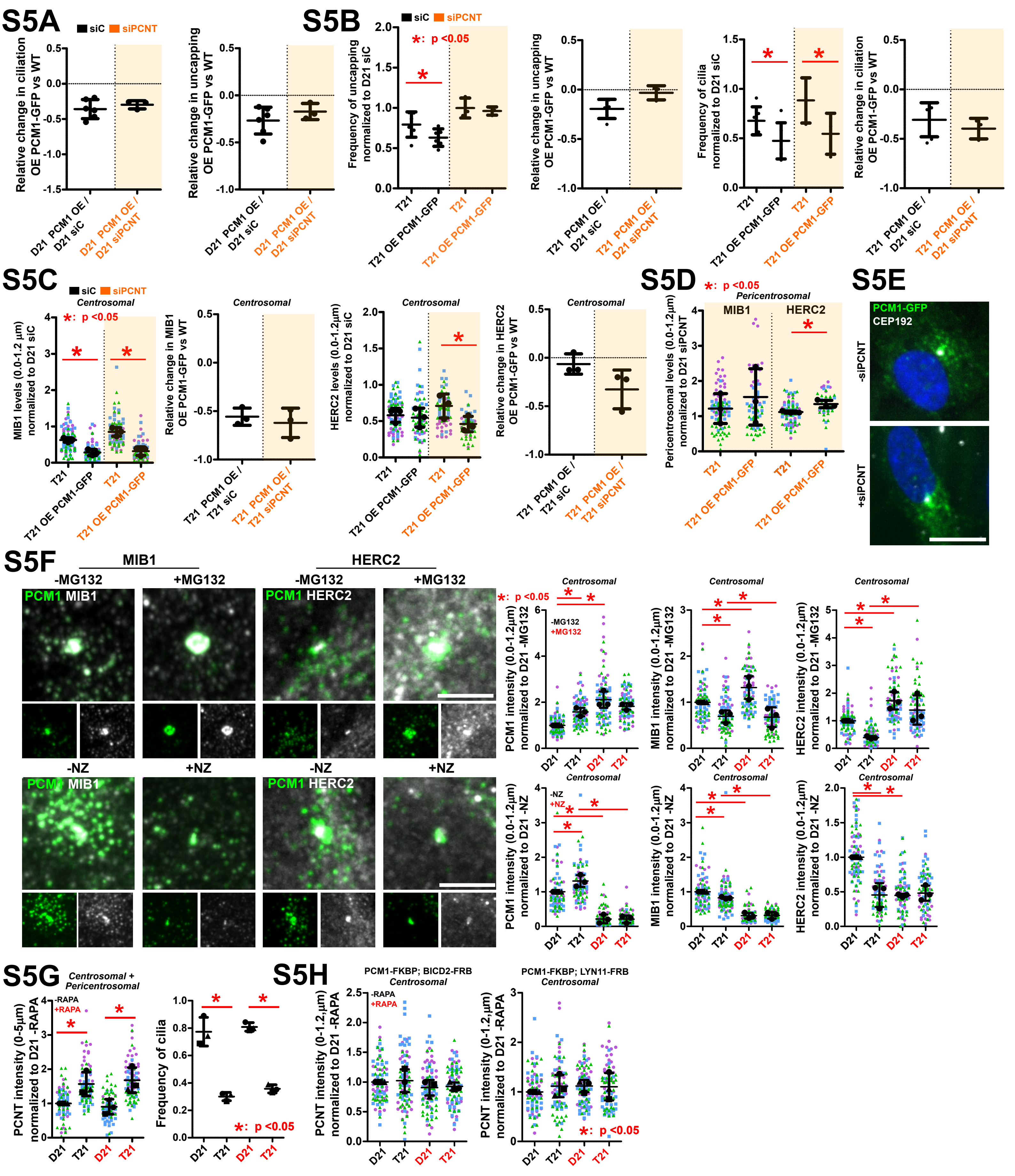
